# Multifaceted genome-wide study identifies novel regulatory loci for body mass index in Indians

**DOI:** 10.1101/670521

**Authors:** Anil K Giri, Gauri Prasad, Khushdeep Bandesh, Vaisak Parekatt, Anubha Mahajan, Priyanka Banerjee, Yasmeen Kauser, Shraddha Chakraborty, Donaka Rajashekar, INDICO, Abhay Sharma, Sandeep Kumar Mathur, Analabha Basu, Mark I McCarthy, Nikhil Tandon, Dwaipayan Bharadwaj

## Abstract

Obesity, a risk factor for various human diseases originates through complex interactions between genes and prevailing environment that varies across populations. Indians exhibit a unique obesity phenotype likely attributed by specific gene pool and environmental factors. Here, we present genome-wide association study (GWAS) of 7,259 Indians to understand the genetic architecture of body mass index (BMI) in the population. Our study revealed novel association of variants in *BAI3* (rs6913677) and *SLC22A11* (rs2078267) at GWAS significance, and of *ZNF45* (rs8100011) with near GWAS significance. As genetic loci may dictate the phenotype through modulation of epigenetic processes, we overlapped discovered genetic signatures with DNA methylation patterns of 236 Indian individuals, and analyzed expression of the candidate genes using publicly available data. The variants in *BAI3* and *SLC22A11* were found to dictate methylation patterns at unique CpGs harboring critical cis- regulatory elements. Further, *BAI3*, *SLC22A11* and *ZNF45* variants were found to overlie repressive chromatin, active enhancer, and active chromatin regions, in that order, in human subcutaneous adipose tissue in ENCODE database. Besides, the identified genomic regions represented potential binding sites for key transcription factors implicated in obesity and/or metabolic disorders. Interestingly, rs8100011 (*ZNF45*) acted as a robust cis-expression quantitative trait locus (cis-eQTL) in subcutaneous adipose tissue in GTEx portal, and *ZNF45* gene expression showed an inverse correlation with BMI in skeletal muscle of Indian subjects. Further, gene-based GWAS analysis revealed *CPS1* and *UPP2* as additional leads regulating BMI in Indians. Our study decodes potential genomic mechanisms underlying obesity phenotype in Indians.

## Introduction

Obesity has become the leading cause for more than 200 medical disorders that affects millions of people worldwide and raises huge economic burden on global health systems [1, 2]. Since 1980, the prevalence of obesity has been doubled in 73 countries including India [3].

Obesity represents a chronic, heterogeneous and complex disorder that precipitates in an individual through cumulative action of genes driven by the environment [4]. Genetic factors contribute nearly 40–70% of inter-individual variability in BMI, a commonly used parameter to assess obesity [4]. So far, genome-wide association studies (GWAS) have identified 227 genetic loci implicated in diverse biological pathways (central nervous system, food sensing, digestion, adipogenesis, insulin signaling, lipid metabolism, muscle/liver biology, and gut microbiome) that may play decisive roles in the development of obese phenotype [5].

GWAS signals for BMI that have been replicated in multiple populations include variants in genes such as *FTO*, *MC4R*, *NEGR1*, *SH2B1*, *TMEM18*, *BDNF*, *FAIM2* and *SEC16B*, besides others [5]. These signals however merely elucidate less than 10% of heterogeneity in BMI presentation, suggesting that a large fraction of genetic determinants remains unknown [5]. One of the reasons for missing heritability can be attributed to population bias in GWAS that have mainly focused on European population, leaving genetic architecture of other populations largely unexplored [6]. Thus, delineating the unknown genetic architecture of remaining global populations may reveal novel population-specific genetic variants, explaining the remaining heritability as well as elucidating the rationale behind population-specific features of obesity.

Indian population is comprised of 4693 diverse communities and various endogamous groups [7]. This genetic diversity was mirrored in our earlier genetic study that attributed new genetic variants within 2q21 region of the human genome for Type 2 diabetes etiology in Indians [8]. Additionally, Indians present distinct features of obesity phenotype that includes greater total, truncal, intra-abdominal, and subcutaneous adipose tissues, as compared to white Caucasians [9]. Moreover, Indians traditionally consume foods enriched in starch and raw sugars, a major contributor to obesity. These reasons suggest that population-specific genetic variants may characterize obesity in Indian population.

We performed a two-phase GWAS in 7,259 Indians (5,973 adults+1,286 adolescents) that uncovered novel GWAS variants in *BAI3* and *SLC22A11* genes and a novel near GWAS significant variant within *ZNF45* gene. Notably, the identified variations in *BAI3*, *SLC22A11*, and *ZNF45* displayed strong associations to various adiposity-related quantitative measures in Indian subjects such as – weight, waist-hip ratio (WHR), waist-circumference (WC), hip-circumference (HC) besides BMI. To expand the functional relevance, we examined DNA methylation marks at the identified loci in peripheral blood in Indians. *BAI3* and *SLC22A11* variants showed strong association with altered DNA methylation patterns at specific CpGs that harbor vital control regions in the genome. Further, analysis of gene regulatory datasets suggested that these variants may distinctively influence key regulatory segments in subcutaneous abdominal adipose tissue. Our study identifies novel loci and functional genetic candidates that may regulate obesity biology in Indians.

### Participants and methods

The study was approved by the Human Ethics Committee of All India Institute of Medical Sciences, New Delhi, India and CSIR-Institute of Genomics and Integrative Biology, New Delhi, India and was conducted in accordance with the principles of Helsinki Declarations.

The adult participants were informed about objectives of study and written consents were taken from all of them. For adolescent study population, prior official permission from school authorities, informed written consent from parents/guardians and verbal consent from participants themselves were obtained before their participation in the study.

### Study Population

The adult participants with more than 18 years of age were Indians of Indo-European origin residing in Northern India and were enrolled through health awareness camps executed in/around Delhi. These participants were also members of INdian Diabetes Consortium (INDICO) [10] and served as normoglycemic controls for Type 2 diabetes (T2D) GWAS executed earlier in lab [8].

Further, adolescent participants of Indo-European origin under the age group of 10-17 years were sampled as part of a GWAS for childhood obesity and related traits in Indians. Participants were enrolled through school health surveys piloted in different zones of Delhi NCR (north, south, east, west, and central regions) as described previously [11–14]. These subjects are well characterized for anthropometric as well as biochemical measurements [11–14].

Blood samples were drawn from participants after overnight fast, and genomic DNA was extracted from peripheral blood using salt precipitation method. Height, weight, waist and hip circumferences (WC and HC) were calculated using standard methods as described earlier [10]. Levels of glucose (FG), HbA1c, insulin (FI), total cholesterol (TC), LDL cholesterol (LDL-C), HDL cholesterol (HDL-C), triglycerides (TG), C-reactive protein (CRP), urea, creatinine, uric acid and blood pressure were measured as described previously [10]. Detailed anthropometric and biochemical characteristics of adult study subjects are presented in Supplementary Table 1. Detailed analysis pipeline employed in study is shown in Supplementary Fig.1.

### Genome-wide association study

#### Discovery Phase

Discovery phase samples were scanned genome-wide using Illumina Human610-Quad BeadChips (Illumina Inc., San Diego, CA) as part of GWAS studies performed previously for T2D and related metabolic traits in our laboratory [8,15–18]. Genotype data was processed through Genome Studio software and further analyzed using PLINK [19]. Samples with missing data for >5% of single nucleotide polymorphisms (SNPs), and with sex-discrepancy between calculated sex and reported sex were removed. We also removed samples with extremely low or high heterozygosity (mean ±3 SD). Related samples with Pi-hat score >0.1875 and potential population outliers (mean ± 6SD) were detected using analysis of first five principal components and were removed. Principal components were calculated using GCTA tool (http://www.complextraitgenomics.com/software/gcta/) [20]. SNPs with minor allele frequency (MAF) <0.01 were excluded from analysis. For SNPs with MAF range between 0.01 and 0.05, those with less than 97% call rate or with Hardy Weinberg Equilibrium (HWE) p<10^−5^ or sex and mitochondrial SNPs were removed.

BMI values were inverse normalized using an inbuilt command in R (http://www.r-project.org/) before association analysis. Association of QC-passed 5,37,246 SNPs with inverse-normalized BMI levels of 1,142 individuals were tested using linear regression model adjusted for age, sex, and first two principal components in PLINK [19]. To find the deviation of p-values, a quantile-quantile (QQ) plot between observed and theoretical p-values was plotted using the qqman package in R [21].

#### Replication Phase

The present study was part of a large genetic study to identify genetic variations regulating various metabolic traits (glycemic, lipids, nitrogen, inflammatory and adiposity parameters) in apparently healthy Indian adults [15–18]. SNPs with discovery phase p◻<◻10^−4^ for any of the considered metabolic traits besides earlier known signals for BMI levels were genotyped in validation phase using GoldenGate technology (Illumina, San Diago, USA). A total of 204 samples (4.22%) were genotyped in duplicates and an error rate <0.01% was detected between technical replicates.

Samples with less than 90% call rate were excluded from the analysis. SNPs with genotype confidence score (confidence value ranging from 0-1, assigned to each genotype call)<0.25, GenTran score (statistical score that mimics evaluations made by human expert’s visual/cognitive systems about clustering behavior of a locus)<0.60, cluster separation score (cluster separation measurement between different genotypes for an SNP that ranges from 0-1)<0.4 and call rate <90% were excluded from analysis. SNPs with minor allele frequency <0.01 and Hardy Weinberg equilibrium (HWE) p<10^−5^ were also removed. After stringent QC, total 4,831 individuals were analyzed in validation phase of the study. BMI values were inverse normalized. Association analysis was carried using linear regression adjusted for age and sex.

We further tested the association of top 3 novel signals (p≤10^−7^) observed in Indian adults during meta-analysis in 1,286 Indian adolescents that were genotyped using Axiom™ Genome-Wide EUR 1 Array. Data for adolescent cohort were also analyzed using standard QC procedures (samples with <90% call rate, SNPs with<95% call rate and HWE p<10^−5^ were excluded) before association. Association analysis was carried using linear regression adjusted for age, sex, and first two principal components. METAL (http://www.sph.umich.edu/csg/abecasis/Metal/) [22] was used for meta-analysis of summary statistics of discovery (adult participants) and validation phases (adult and adolescent participants) of study using fixed-effect inverse variance method.

Moreover, we performed an *in-silico* replication of identified SNPs in 2,078 South Asian subjects from United Kingdom Biobank (UKBB) (http://biobank.ctsu.ox.ac.uk) [23]. Further earlier association status of discovered variants and genes were retrieved from Type 2 diabetes knowledge portal [24].

### Statistical Power of study

Power of study was calculated using Quanto software (http://biostats.usc.edu/Quanto.html) assuming an additive genetic model for allele frequencies ranging from 0.001-0.5. For power calculations, two-tailed test at significance level of 0.05 with effect size ranging from 0.1-0.5, obtained from literature, was used. The average BMI was taken as 25.72 kg/m^2^ with a standard deviation of 5.15 kg/m ^2^ for power calculation.

### Combined risk score analysis

To identify the cumulative effect of 14 established and 3 novel SNPs on BMI levels in Indians, we performed allele dosage analysis by classifying the subjects on the basis of number of “effective” risk alleles as described earlier [25]. The analysis involved samples in which genotypes at all 17 SNPs were available. We calculated effective unweighted as well as weighted allele dosage score (ADS) for this purpose.

An unweighted ADS was computed as sum of number of risk alleles for all 17 SNPs per individual. However, weighted ADS was calculated as the weighted mean of the proportion of risk alleles at 17 SNPs (i.e. 1 for two risk alleles, 0.5 for one risk allele, and 0 for no risk allele) with weights as the relative effect sizes of different SNPs. The “effective” number of risk alleles was derived by multiplying weighted ADS by 34 (maximum number of risk alleles corresponding to 17 SNPs).

BMI values were inverse normalized. Effect sizes (beta) and P-values for overall trend in total subjects was calculated using linear regression analysis in SPSS version 25.0 (https://www.ibm.com/in-en/analytics/spss-statistics-software) to identify change in BMI levels with every unit increase in number of “effective” risk alleles. For unweighted and weighted risk score analysis, subjects with <10 and <80 number of “effective” risk alleles respectively were taken as the reference group to calculate risk of obesity for different risk groups. For this, subjects were classified into two groups – subjects with BMI <25 kg/m2 (normal weight) and subjects with BMI ≥ 25 kg/m2 (overweight/obese). The p-values and odds ratio while comparing the different groups for risk of obesity was calculated using chi-squared test statistic.

### Imputation Analysis

Imputation analysis of *BAI3* and *ZNF45* loci (signals with discovery p<0.05) was performed in GWAS dataset as described previously [8]. For reference panel, 1000 Genomes Phase 3 population was used. In brief, pre-phasing for respective chromosomes was carried out using SHAPEIT [26]. A region of 1 Mb on either side encompassing the LD block of the variant was imputed using IMPUTE2 [27]. Stringent QC was performed on imputed SNPs that followed: Certainty ≥ 0.90, Info ≥ 0.5 and MAF ≥ 0.01 (Supplementary Fig. 2). Imputed SNPs that passed QC were tested for association with inverse normalized BMI levels in Indians using PLINK [19]. Age, sex, and the first two principal components were employed as covariates in the model.

### Correlation among adiposity traits

Correlations among adiposity measures – BMI, weight, WC, HC and WHR were calculated using corrplot package in R (http://www.r-project.org/). R corrplot function was applied to plot the correlation matrix. Since adiposity measures were strongly correlated, association of SNPs in *BAI3*, *SLC22A11*, and *ZNF45* were also tested with other adiposity traits - weight, WHR,WC, HC and related metabolic traits - FG, HbA1c, FI,TC, LDL-C, HDL-C, TG, CRP, urea, creatinine, uric acid, blood pressure (systolic-SBP and diastolic-DBP). Covariates for adiposity traits association analysis were age, sex, and the first two principal components in discovery phase and age, sex in validation phase while BMI was additional covariate for association analysis with metabolic traits in both phases of adult Indians using linear regression based model in PLINK [19]. METAL (http://www.sph.umich.edu/csg/abecasis/Metal/) [22] was used for meta-analysis of discovery and validation phases of SNP-trait associations using fixed-effect inverse variance method.

### DNA methylation analysis

We performed whole-genome DNA methylation experiments in peripheral blood of 236 Indian subjects used in discovery phase using Infinium HumanMethylation450K BeadChip. Methylation data was analyzed through ENmix and Minfi packages in R with BMIQ normalization [28–30] as described previously [31]. For meth-QTL analysis, we selected CpGs present in SNP related genes to figure out any alterations in methylation level of nearby CpGs due to presence of identified SNPs.

Sample quality control involved removal of samples with sex discrepancy, samples that failed during bisulphate conversion (samples with intensity 3 SD away from mean intensity for C1, C2, C3 and C4 probes), and samples with more than 5% CpG sites missing. CpG quality control include removal of CpGs with bead count less than 3 in 5% of samples, detection p-value >0.01 for less than 1% of samples. Additionally, CpGs falling in sex chromosomes, cross-hybridization probes and polymorphic CpGs were excluded from analysis [32]. CpGs with 100% call rate in all the samples were carried forward for meth-QTL analysis. CpG outliers were fixed using fixMeth-Outliers command in Minfi [29]. Further, data was regressed for confounders such as – cell composition, age, sex, BMI, bisulphate conversion efficiency and plate number. Methylation values for 53 CpGs (19 CpGs present in *BAI3*, 25 CpGs in *SLC22A11* and 9 CpGs in *ZNF45*) were extracted, and SNP-CpG association analysis was executed using linear regression model in PLINK [19].

### Integration of gene regulatory data

GTEx-portal-v7 (The Broad-institute of MIT and Harvard) [33] was used for retrieving Global expression-QTL (eQTL) and tissue expression profiling data for identified SNPs and genes respectively. Whole Genome Bisulphite Sequencing data (WGBS) for adipose tissue was obtained from female and male adult subjects aged 30 and 34 years respectively from ENCODE dataset [34]. Human ATAC-seq data of subcutaneous adipose tissue was derived from an adult female aged 53 years from ENCODE [34].

ChIP-Seq data for regulatory histone marks belong to subcutaneous adipose tissue of five adult females aged 25, 41, 49, 59 and 81 years, and was acquired from ENCODE [34]. ENCODE data for DNase I hypersensitive sites in 95 cell types was also examined. Experimentally defined as well as predicted transcription factor (TF) binding sites were obtained from ENCODE and JASPAR database respectively [34,35]. Additionally, likely chromatin interaction potential data was retrieved from GeneHancer database that features human regulatory elements (enhancers and promoters) and their inferred target genes [36]. UCSC genome browser was used to schematically visualize gene regulatory aspects of discovered regions [37].

### Expression analysis of identified genes

We examined correlation between BMI levels and expression level of discovered genes in adipose and skeletal muscle tissue of 6 Indian subjects (3 males + 3 females) using our earlier published data generated through Illumina HumanHT-12 v3 Expression BeadChip arrays [38]. Each sample was assessed 4 times to reduce technical variability in gene expression data. Pearson correlation was calculated between average of expression values for respective gene probes and BMI.

### Gene-based association study

Besides SNP based GWAS analysis, we also conducted a univariate gene-based genome-wide association scan using an effective chi-squared method (ECS) employed in knowledge-based mining system for genome-wide genetic studies (KGG v4) accessible at http://statgenpro.psychiatry.hku.hk/limx/kgg/download.php. Gene-based association study may identify novel gene sets for a population based on associated marker buildup in whole genes. Genome–wide markers with association p-values for BMI were used as input for KGG v4.

As *BAI3*, *SLC22A11*, and *ZNF45* signals were robustly linked to a majority of adiposity parameters besides BMI (weight, WHR, WC, HC), we also used a multivariate gene-based association test for these genomic regions utilizing extended Simes method (MGAS) [39]. Multivariate approach provides gene-based testing of several correlated phenotypes in large number of unrelated subjects. Association p values of discovery phase SNPs within 2 Mb region of *BAI3*, *SLC22A11*, and *ZNF45* for all adiposity parameters and correlation values among these traits were provided in MGAS model using KGG v4.

1000 Genome phase III data of European, African, American, East Asian, and South Asian populations were used for computing Linkage disequilibrium (LD) among all the markers within the respective region.

## Results

The present study was effectively powered to detect variants with similar effect sizes as observed in earlier GWAS studies for BMI in literature (>98%) (Supplementary Fig.3). Under null hypothesis, QQ plot displayed a good concordance (Supplementary Fig.4). The genomic inflation factor (λ) was estimated to be 1.06 signifying homogeneity of study population.

### Genome-wide association analysis of BMI

Meta-analysis of summary results from discovery and validation phases (N=7,259) revealed *BAI3* (rs6913677, P = 1.08 × 10^−8^) and *SLC22A11* (rs2078267, 4.62 × 10^−8^) as novel GWAS signals for BMI in Indians (Table 1, Fig.1). A novel near GWAS level association for BMI in *ZNF45* loci (rs8100011, P=1.04 ×10^−7^) was also observed (Table 1, Fig.1).

**Table 1.** Association status of SNPs with BMI. Effect sizes were calculated with respect to risk alleles. Association results presented here have been obtained from meta-analysis of summary results from discovery and validation phases in Indian adult and adolescent cohort. Dir: direction; Het-P: p-value for heterogeneity in effect sizes in the meta-analysis; I^2^: Chi-square value for heterogeneity test. Direction ++/−− represents a concordance between the discovery and replication phase. Proxy SNPs - rs11752858 and rs2277312 have been utilized for rs6913677 and rs2078267 respectively for analysis in the adolescent cohort. rs2078267 was selected for genotyping in validation phase because it was associated with waist circumference and serum uric acid level at a p-value less than 10^−4^

|  |  |  | Discovery phase in<br>adults<br>(N = 1,142) |  | Validation phase in<br>adults<br>(N = 4,831) |  | Validation phase in<br>adolescents<br>(N =1,286) |  | Meta-analysis |  |  |  |
| --- | --- | --- | --- | --- | --- | --- | --- | --- | --- | --- | --- | --- |
| Trait | SNP<br>(RA/OA)<br>Gene | RAF | BETA<br>(SE) | P | BETA<br>(SE) | P | BETA<br>(SE) | P | BETA<br>(SE) | P | Het-P<br>(I<sup>2</sup>) | Dir |
| BMI | rs6913677<br>(G/A)<br>*BAI3* | 0.57 | 0.71<br>(0.19) | 1.21 x10<sup>-4</sup> | 0.38<br>(0.11) | 3.46 x10<sup>-4</sup> | 0.55<br>(0.20) | 5.70 x10<sup>-3</sup> | 0.47<br>(0.08) | 1.08x10<sup>-8</sup> | 0.26<br>(2.87) | +++ |
|  | rs2078267<br>(A/G)<br>*SLC22A11* | 0.41 | 0.16<br>(0.19) | 0.40 | 0.52<br>(0.10) | 6.76 x10<sup>-7</sup> | 0.53<br>(0.20) | 7.12 x10<sup>-3</sup> | 0.45<br>(0.08) | 4.62x10<sup>-8</sup> | 0.23<br>(23.78) | +++ |
|  | rs8100011<br>(A/G)<br>*ZNF45* | 0.55 | 0.64<br>(0.19) | 7.17 x10<sup>-4</sup> | 0.40<br>(0.11) | 1.76 x10<sup>-4</sup> | 0.40<br>(0.20) | 0.05 | 0.44<br>(0.08) | 1.04 x10<sup>-7</sup> | 0.51<br>(1.33) | +++ |

**Fig. 1.**
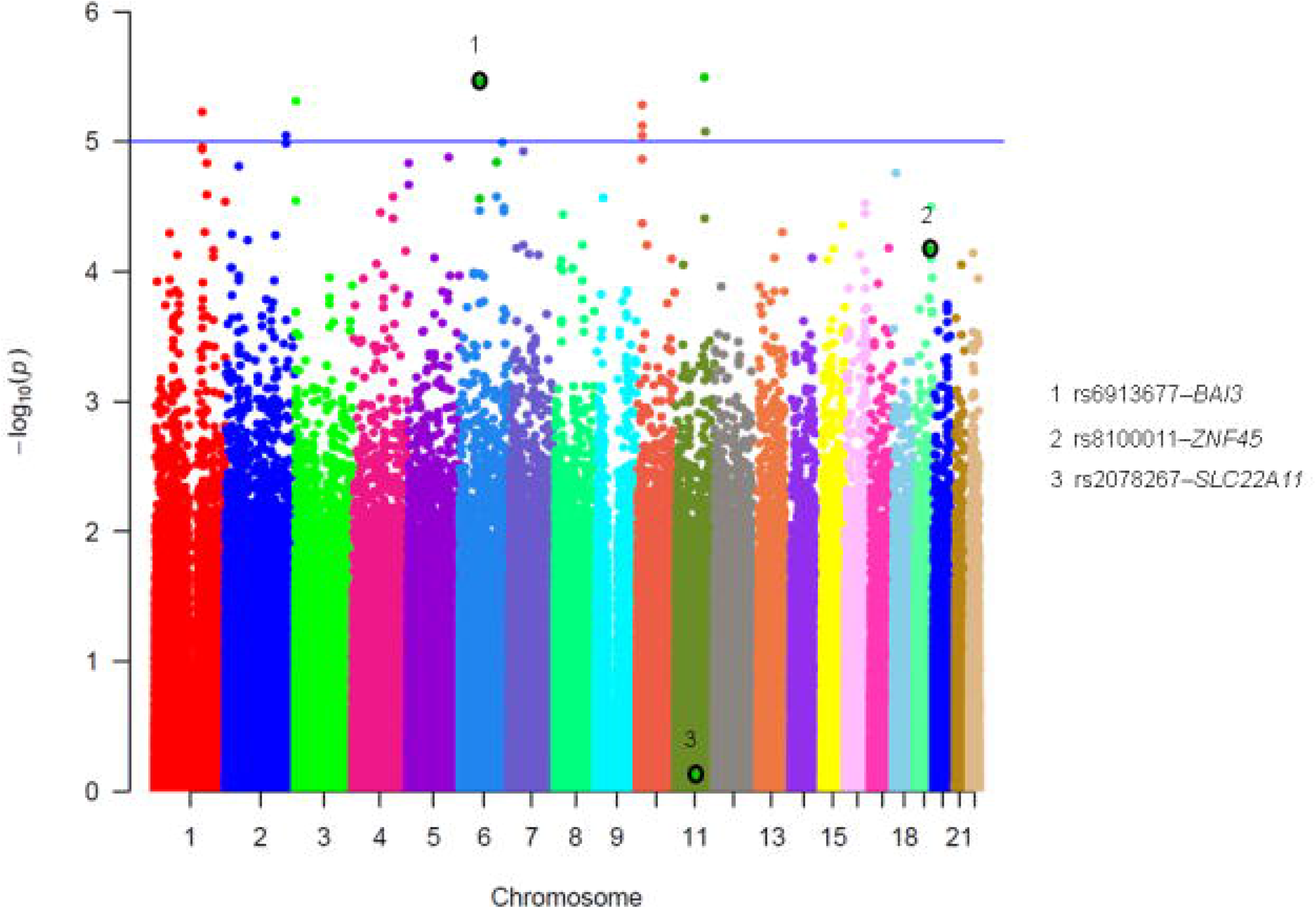
Manhattan plot for the SNPs associated with BMI in discovery phase. The −log10 p-values for the association of genotyped SNPs are plotted as function of genomic position (National Center for Biotechnology Information Build 37). The p-value was calculated using logistic regression adjusted for age, sex, PC1, and PC2 in discovery phase analysis. Each chromosome (CHR) has been represented with a unique color.

**Fig.2.**
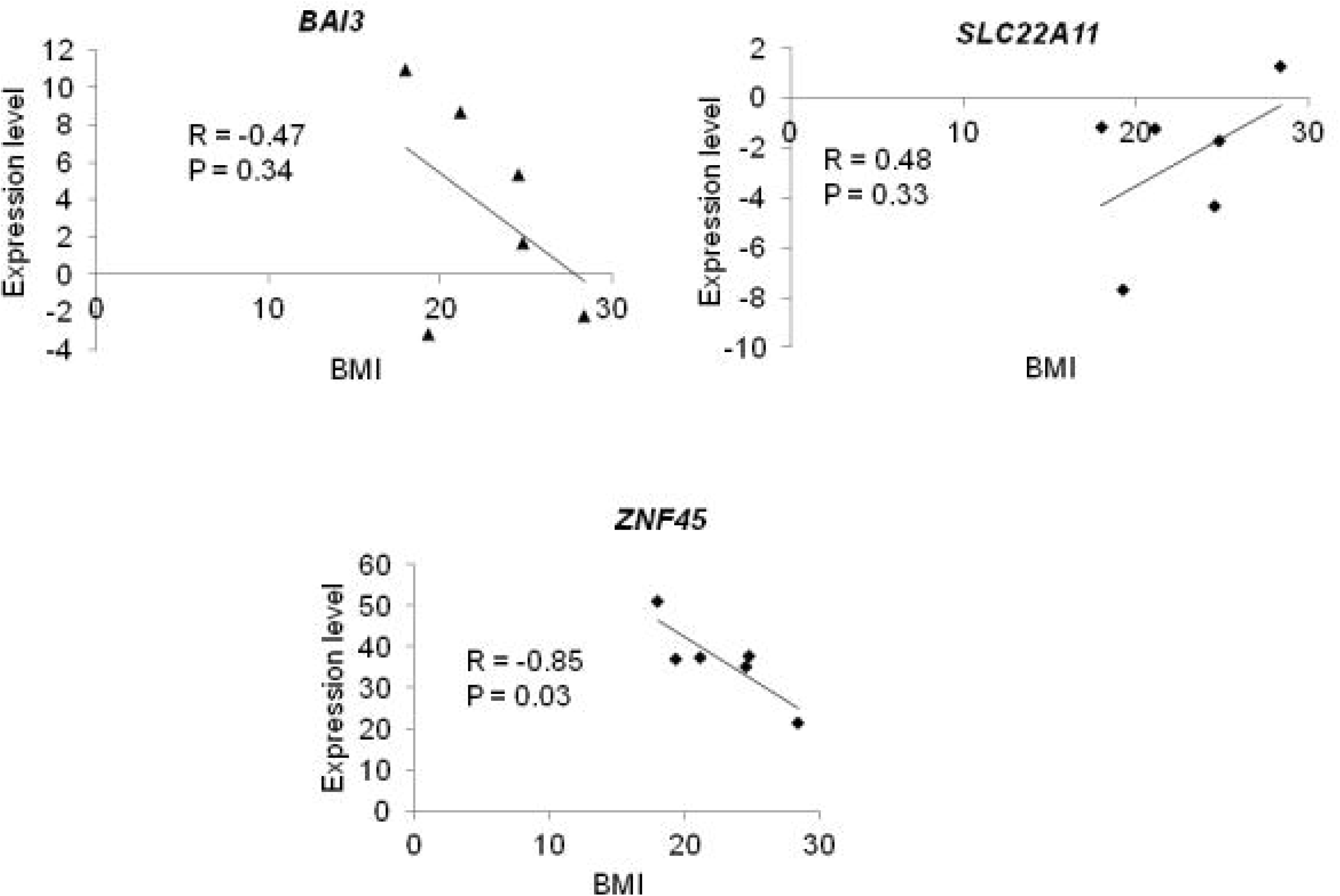
Correlation between expressions of identified genes in skeletal muscle and BMI in Indians. Gene expression of *BAI3*, *SLC22A11* and *ZNF45* in skeletal muscle tissue of 6 Indian subjects (3 males and 3 females) has been correlated with BMI levels of study subjects. Pearson correlation was calculated between average of expression values for respective gene probes and BMI.

Additionally, among the known variants for BMI that were genotyped in replication phase in adults (N=4,831), previously reported signals near *MC4R (*rs17782313 and rs12970134) showed replication with GWAS significance for BMI levels (Supplementary Table 2). Moreover, variants in/near *BDNF*, *FTO*, *LOC199899*, *MTNR1B*, *TCF7L2*, *FADS1*, *KCTD15*, *TMEM18* and *SH2B1* replicated with nominal significance for BMI (Supplementary Table 2).

### Allele dosage analysis

We analyzed the combined effect of multiple alleles at identified loci - 14 established (as listed in Supplementary Table 2) and 3 novel loci (rs6913677-*BAI3*, rs2078267-*SLC22A11* and rs8100011-*ZNF45*) on BMI levels using weighted and unweighted allele dosage analysis. Results suggested a significantly increased levels of BMI by 0.15 units with the rise in each unit of “effective” risk allele (P = 4.23×10^−21^) (Supplementary Fig.5a). Subjects with >172 “effective” risk alleles (2%) displayed 3.1 times enhanced risk for overweight/obesity in comparison to subjects having less than 80 “effective” risk alleles (11%) (P = 2.1×10^−5^) (Supplementary Fig.5a). An unweighted allele dosage analysis revealed that subjects with 25-29 “effective” risk alleles (2.33%) displayed 5-fold enhanced risk for overweight/obesity in contrast to subjects having less than 10 “effective” risk alleles (1.94%) (P = 9.35×10^−7^) (Supplementary Fig.5b).

### *BAI3*, *SLC22A11* and *ZNF45* variants are novel loci for BMI

We examined former reported associations within *BAI3*, *SLC22A11*, and *ZNF45* for BMI levels (Supplementary Fig.6 (a, b and c)). The strongest reported associations for BMI included the SNPs - rs618714 (p=2.10×10^−7^), rs693591 (p=0.04) and rs454376 (p=0.01), within *BAI3*, *SLC22A11*, and *ZNF45* respectively in GIANT UK Biobank GWAS comprising majorly of Europeans. Interestingly, recently a *BAI3* variant - rs513357 displayed genome-wide significance (p=8×10^−9^) with BMI in a multi-ethnic cohort that comprised of European, Hispanic/Latin American, East Asian, African American/Afro-Caribbean, Africans and South Asians [40]. Our identified *BAI3* variant rs6913677 was in weak LD with this earlier observed rs513357 loci in Indians (r2=0.16 - our data), South-Asians (r2=0.10), Africans (r2=0.002), Americans (r2=0.02), East-Asians (r2=0.038) and Europeans (r2=0.13) (https://ldlink.nci.nih.gov/?tab=ldmatrix) [41]. Further, the presently discovered variants for BMI in Indians (rs6913677, rs2078267 and rs8100011) were only nominally attributed for BMI and related traits in earlier studies (Supplementary Table 3).

### *In-silico* replication of novel variants

*In-silico* replication of novel BMI signals in South Asian individuals from UK BioBank did not improve the association significance for any of the three loci. It however revealed similar direction of BMI association for *SLC22A11* (rs2078267), and *ZNF45* (rs8100011) (Supplementary Table 4).

### Gene expression correlation analysis

Expression correlation analysis between BMI levels of Indian subjects and gene expression levels of *BAI3*, *SLC22A11*, and *ZNF45* in adipose and skeletal muscle tissues revealed a strong negative correlation between *ZNF45* gene expression and BMI (R= − 0.85, P = 0.03) in skeletal muscle of study subjects as shown in Fig.2. Further, publicly available data mining in GTEx portal revealed *ZNF45* variant - rs8100011 as strong *cis*-eQTL in subcutaneous adipose, thyroid, skin, and tibial nerve (Supplementary Table 5, Supplementary Fig.7). The eQTL results from GTEx domain suggested that the double risk allele genotype for BMI (“Genotype AA”; rs8100011) was associated with reduced expression of *ZNF45* in various human tissues (Supplementary Fig.7). This is consistent with our observations of increase in BMI with decrease in expression of *ZNF45* (facilitated by risk genotype AA of rs8100011) in skeletal muscle of obese subjects (Fig.2).

Further, expression correlation analysis for *BAI3* and *SLC22A11* exhibited modest negative (R= − 0.47, P = 0.34) and positive correlation (R= 0.48, P = 0.33) respectively with BMI levels in skeletal muscle of Indian subjects (Fig. 2).

### Imputation analysis of novel regions

For *BAI3*, we observed 7 variants that showed marginally greater significance for association with BMI levels than index SNP rs6913677 (Supplementary Table 6a). At *ZNF45* loci, 6 variants other than the index SNP rs8100011 showed slightly greater significance for association with BMI levels (Supplementary Table 6b). Few of these imputed variants (rs10945151, rs55736013, and rs11880216) overlapped with binding sites for key transcription factors such as PBX1, HOXC9, KLF4, KLF5, KLF9, and IRF3, which are known for their involvement in adipogenesis related processes. Further, imputed variants of *ZNF45* (rs55736013 and rs11880216)were also observed as robust cis-eQTLs in human subcutaneous adipose tissue, thyroid, skin, and tibial nerve, similar to the genotyped *ZNF45* variant, in the GTEx database [33].

### *BAI3*, *SLC22A11* and *ZNF45* loci are shared genetic determinants for adiposity traits in Indians

Adiposity traits (BMI, weight, WHR, WC and HC) were observed to effectively correlate with each other in Indians (Supplementary Fig.8). BMI levels were observed to strongly correlate with weight (correlation coefficient: 0.83), WC (correlation coefficient: 0.75), HC (correlation coefficient: 0.85) and poorly with WHR (correlation coefficient: 0.07). Weight correlated strongly with WC (correlation coefficient: 0.83), HC (correlation coefficient: 0.72) and modestly with WHR (correlation coefficient: 0.33). WHR showed modest positive correlation with WC (correlation coefficient: 0.57) and negative correlation with HC (correlation coefficient: −0.17). WC showed strong positive correlation with HC (correlation coefficient: 0.71).

Accordingly, we looked at association status of *BAI3*, *SLC22A11*, and *ZNF45* with all of these adiposity and related metabolic measures. Interestingly, *BAI3* variant (rs6913677) showed robust association with WC (p-value=8.75◻×◻10^−7^), HC (p-value=4.90◻×◻10^−7^), weight (p-value=2.60◻×◻10^−7^), LDL-C (1.73◻×◻10^−4^), TC (1.95◻×◻10^−3^) and SBP (p-value=0.03) besides BMI in Indians (Supplementary Fig.9, Supplementary Table 7). Further, *SLC22A11* variant (rs2078267) showed strong association with WC (p-value=1.23◻×◻10^−5^), HC (p-value=2.30◻×◻10^−6^), weight (p-value=2.25◻×◻10^−5^) and uric acid levels (p-value=3.26◻×◻10^−11^) (Supplementary Fig.9, Supplementary Table 7). Also, *ZNF45* variant (rs8100011) displayed strong association with WHR (p-value=5.04◻×◻10^−3^), WC (p-value=6.78◻×◻10^−8^), HC (p-value=1.20◻×◻10^−6^), and weight (p-value=3.70◻×◻10^−9^) (Supplementary Fig.9, Supplementary Table 7). These results suggest that identified novel signals influence the levels of various adiposity features and related metabolic traits in Indians.

### *BAI3*, *SLC22A11*, and *ZNF45* loci are regulatory variations

These genes were observed to be moderately expressed in human subcutaneous and visceral adipose tissues with reasonable ubiquitous expression in liver, skeletal muscle, tibial nerve, pancreas, thyroid, and whole blood. (Supplementary Fig.10). In order to understand functional impacts of identified variants, we first examined their open chromatin features, regulatory histone marks and DNA methylation patterns in human subcutaneous adipose tissue. Besides, we examined predicted TFs that bind to DNA regions of identified variants and their chromatin interaction potential.

*BAI3* variant (rs6913677) represented repressive chromatin assisted by weak ATAC-seq signals in subcutaneous adipose tissue, and absence of DNase I hypersensitivity clusters in multiple human cell-types (Supplementary Fig.11a). This was further supported by enrichment of heterochromatin-specific histone marks H3K27me3 and H3K9me3 around rs6913677 (Supplementary Fig.11a). Interestingly, rs6913677 displayed strong predicted binding sites for several crucial transcription factors implicated in obesity - PPAR-α, PPAR-γ, E2F4, HNF4G, ZNF263 and SP2 (Supplementary Fig.11b) with highly conserved DNA binding motifs (denoted by high TF bit-score) for these key transcription factors (Supplementary Fig.11c). In addition, WGBS data revealed differential methylation at rs6913677 position in adipose tissue of male and female samples, wherein 50% of sequence reads were found to be methylated in male sample with no methylation in female sample, indicating sex-specific regulatory potential of this variant (Supplementary Fig.11b).

In contrast, *SLC22A11* variant (rs2078267) constituted strong open chromatin characteristics supported with robust ATAC-seq and DNase I hypersensitivity signals in subcutaneous adipose tissue and various other cell types respectively (Supplementary Fig.12a). Strong peaks of H3K4me1 accompanied by feeble peaks of H3K27ac and H3K4me3 around rs2078267 in subcutaneous adipose tissue represent an active enhancer region (Supplementary Fig.12a). This was supported by GeneHancer database that demonstrated strong enhancer functionality of rs2078267 variant targeting *TRMT112*, *CDC42BPG* and *SF1* genes in its vicinity (Supplementary Fig.12b). Enhancer ability shows ‘double elite’ status that reflects a greater likelihood of prediction accuracy for both enhancer and target gene. Interestingly, variant rs2078267 displays strong binding sites for experimentally identified potential obesity and metabolic disease associated transcription factors. These include - NFIC, KAP1, TCF7L2, ZNF263, GTF2F1, EP300, POU5F1, FOXA1, TBP, RXRA, JUND, MYC, BHLHE40, SP1, ARID3A, RCOR1, HDAC2, FOXA2, CHD2, USF1, ATF1, GATA3, JUN, MAZ, BRCA1, FOXP2, ATF3, CEBPD, FOS, CEBPB, FOSL2 and USF2 (Supplementary Fig.12b). Further, none of sequenced reads were found to be methylated in adipose tissue around variant rs2078267 in WGBS dataset (Supplementary Fig.12b).

*ZNF45* variant (rs8100011) exhibited extensive enrichment of active H3K36me3 and H3K4me3 marks and feeble peaks of repressive H3K27me3 and H3K9me3 marks in subcutaneous adipose tissue, which is a signature for active gene transcription (Supplementary Fig.13a). The variant region indicated strong chromatin interaction potential of *ZNF45* with neighboring genes – *ZNF222*, *ZNF230*, *ZNF283* and *ZNF221* (Supplementary Fig.13b).

### Methylation quantitative trait loci (meth-QTL) analysis

To investigate likely functional roles of identified variants, we overlapped genetic data with DNA methylation data generated from peripheral blood in Indians, and performed meth-QTL analysis that identified associated BMI variants to regulate methylation levels of nearby CpG sites.

Meth-QTL analysis revealed rs6913677 as *cis*-methylation-QTL for a CpG, cg17094144, in *BAI3* gene in peripheral blood in Indians (Table 2). Additionally, rs2078267 displayed robust association with differential methylation patterns for 5 CpGs within *SLC22A11* gene (*cis*-methylation-QTL) in blood tissue in Indians (Table 2).

**Table 2.** Meth-QTL analysis for associated signals in 236 adult subjects who have been genotyped in discovery phase of study. P-value has been obtained from association of SNPs with methylation level at corresponding CpG sites using PLINK. CpG sites have been annotated using Illumina 450K BeadChip manifest file. CHR: Chromosome; BP: Base position; CI: Confidence interval. Signals with p-value<0.05 have only been shown.

| SNP |  |  |  |  | CpG |  |  |  |  |  |
| --- | --- | --- | --- | --- | --- | --- | --- | --- | --- | --- |
| Name | CHR | BP | Gene | A1/A2 | Name | CHR | BP | Gene | BETA (95% CI) | P |
| rs6913677 | 6 | 69860183 | *BAI3* | G/A | cg17094144 | 6 | 69345881 | *BAI3* | 0.01 (0.00 – 0.01) | 0.03 |
| rs2078267 | 11 | 64090690 | *SLC22A11* | A/G | cg00270878 | 11 | 64334216 | *SLC22A11* | -0.03 (-0.04 - -0.01) | 4.44x10<sup>-4</sup> |
| rs2078267 | 11 | 64090690 | *SLC22A11* | A/G | cg07086353 | 11 | 64257659 | *SLC22A11* | -0.01 (-0.02 - 0.00) | 2.20x10<sup>-3</sup> |
| rs2078267 | 11 | 64090690 | *SLC22A11* | A/G | cg08822897 | 11 | 64258102 | *SLC22A11* | -0.05 (-0.07 - -0.04) | 4.96x10<sup>-10</sup> |
| rs2078267 | 11 | 64090690 | *SLC22A11* | A/G | cg09337943 | 11 | 64335732 | *SLC22A11* | 0.01 (0.00 - 0.01) | 4.09x10<sup>-3</sup> |
| rs2078267 | 11 | 64090690 | *SLC22A11* | A/G | cg18158438 | 11 | 64322993 | *SLC22A11* | -0.01 (-0.01 - 0.00) | 0.01 |

Since the identified genes are nominally expressed in human blood (Supplementary Fig.14), we retrieved gene regulatory features for associated CpG sites from K562 leukemia cell line. Associated CpG sites for meth-QTL - rs6913677 is located in exon 1 and 5’ UTR of *BAI3* that may serve as a strong promoter with enrichment of binding sites for RBBP5,EZH2,MYC, RFX5 and MAX transcription factors (Supplementary Table 8). Similarly, meth-QTL - rs2078267 (*SLC22A11*) was robustly associated with five CpG sites that resided in important functional regions - gene body, TSS200 (200 bases upstream from transcription start site) or were intergenic with enhancers or insulators specific functions (Supplementary Table 8).

However, we did not observe rs8100011 (*ZNF45*) as meth-QTL in peripheral blood in Indians even though *ZNF45* is moderately expressed in blood (Supplementary Fig.14). We also examined the WGBS dataset around identified variants in classical monocytes (CD14-positive), B-cells and T-cells from ENCODE dataset. All these blood cells showed lack of any DNA methylation mark at rs8100011 (*ZNF45*) loci in contrast to rs6913677 (*BAI3*) and rs2078267 (*SLC22A11*) (Supplementary Fig.15). Similarly, no methylation status for rs8100011 (*ZNF45)* was observed in adipose tissue derived from ENCODE dataset (Supplementary Fig.13b).

### Gene-based GWAS analysis

As SNPs alone cannot fully explain the heritability of complex traits, we also implemented a gene-based GWAS analysis to identify complementary genetic determinants for BMI in Indians. The analysis uncovered distinct novel loci that were not detected earlier in SNP based association method. Protein-coding genes *CPS1* and *UPP2* were strongest signals (p-value≤1×10^−8^) associated with BMI for the first time (Supplementary Table 9). This was followed by additional protein-coding genes *ACOXL*, *FAM71E2*, *NRCAM*, *SLC25A12*, *PKD1L3*, *UBA5* and *APBA2*, and non-coding RNA genes *LINC00358, LINC01142* and *SLC16A12-AS1* that associated modestly (p-value≤1×10^−5^) with BMI in Indians (Supplementary Table 9). Further, in agreement with SNP-based analysis, *BAI3* and *ZNF45* genes also persisted significance in gene-based analysis (respective p= 0.02 and 3.55×10^−5^).

Since *BAI3*, *SLC22A11* and *ZNF45* loci associated robustly with most adiposity traits in Indians, we also implemented a multivariate gene-based testing for all three loci using extended Simes method (MGAS) employing gene-based testing for various correlated phenotypes in independent samples. The gene-based analysis suggested *BAI3* and *ZNF45* as leading candidates within their respective 2 Mb genomic regions (p-value=0.01 and 5.50×10^−4^ respectively) associated with all adiposity measures in Indians (Supplementary Table 10).

## Discussion

Present study investigated genome-wide signatures regulating BMI in Indians. We discovered novel GWAS significant loci in – *BAI3* (brain-specific angiogenesis inhibitor) and *SLC22A11* (solute carrier family 22 member 11) followed by *ZNF45* (zinc finger protein 45) locus with near GWAS significance.

Our study uncovered *BAI3* that belongs to cell-adhesion class of seven-span transmembrane G protein-coupled receptors as regulating adiposity traits in Indians [42]. *BAI3* is widely expressed in different regions of human brain (cerebral hemisphere, cerebellum, and pituitary) [33]. One of its 4 ligands, C1ql4 (Complement C1q-Like Protein 4) has been identified as a negative regulator of adipogenesis in mouse model via inhibiting p42/44-Mitogen-Activated Protein Kinase signaling pathway (p42/44-MAPK) [43, 44]. This pathway plays a critical role in regulating the balance between adipocyte growth and differentiation [45]. Interestingly, over-expression of C1ql4 repressed the differentiation of 3T3-L1 adipocytes marked up by parallel reduction in transcript as well as protein levels of PPAR-γ and C/EBP-α, major transcription factors that drive adipogenesis [43].

C1ql4 protein is a small secreted signaling molecule expressed variably in human brain [33], wherein globular C1q domains bind strongly with extracellular thrombospondin-repeat domain of BAI3 receptor and may direct synapse formation and maintenance in human brain [44]. Owing to consideration that both *BAI3* and *C1ql4* genes are also nominally expressed in human subcutaneous and visceral adipose tissues [33], we speculate that BAI3 receptor can mediate the inhibitory effect of C1ql4 ligands on adipocyte differentiation plausibly via p42/44-MAPK pathway.

A modest enrichment for gene repression histone marks (H3K27me3 and H3K9me3) surrounding *BAI3* locus in human subcutaneous adipose tissue designates it a facultative heterochromatin zone. We observed that alternate alleles of *BAI3* variant rs6913677 significantly influenced the methylation levels of CpG site cg17094144 in Indians. The CpG site resided within active promoter region of *BAI3*, and is surrounded by regulatory histone modifications with definite binding sites for critical transcription factors. Altered DNA methylation can modulate the binding of these TFs and methyl CpG associated proteins, thereby influencing gene expression. For instance, DNA methylation in intron 1 of *HIF3A* presents robust association with BMI levels and significant negative correlation with *HIF3A* mRNA levels in adipose tissue [46]. Additionally, DNA methylation marks in blood at four CpG sites residing in *LGALS3BP*, *RORC*, *SOCS3* and *ANGPT4* retained robust association with BMI levels in American Women [47].

Identification of *BAI3* variant rs6913677 as cis-meth-QTL demonstrates that variant may fine-tune adiposity via affecting *BAI3* gene expression. In addition, this variant functions as strong binding motif for known TFs like PPAR-α, PPAR-γ, E2F4, HNF4G, ZNF263 and SP2, known to play a major role in obesity [35]. Likewise, imputed variant of *BAI3* - rs10945151 also marked a strong seat for adiposity related TFs such as PBX1 and HOXC9 [35]. PBX1 controls the process of adipogenesis in stage-dependent manner, in both mouse and human [48]. Additionally, HOXC9 is involved in adverse fat distribution, metabolism, adipose tissue function and development of obesity [49]. Alternate alleles of *BAI3* variant can efficiently influence the binding of these TFs to their respective binding motifs and thereby alter the *BAI3* gene expression levels. Variable gene expression may subsequently modify levels of expressed *BAI3* protein that will influence downstream receptor-ligand interactions and consequently adipocyte differentiation in adipose tissue.

Intriguingly, multiple intergenic GWAS loci for BMI near *BAI3* were recently observed in GIANT UK Biobank GWAS conducted in ~700,000 individuals of European ancestry [50]. Till now, only one variant - rs513357 within intron 2 of *BAI3* gene presented GWAS level association with BMI in a large multi-ethnic cohort [40]. The reported variant displayed weak LD with our discovered *BAI3* variant - rs6913677 (intron 16) as well as with our imputed *BAI3* variants in Indians and several other populations [41]. Differences in allele frequencies and haplotype structures across diverse human populations may attribute for these population-specific associations. Further, a few GWAS variants within/near*BAI3* have also been identified for omega-6 fatty acid, namely linoleic acid and arachidonic acid levels in European individuals [24] as well as for triglycerides levels in Native American population [51]. Besides, a large number of related metabolic traits–waist circumference, waist-hip ratio, dihomo-gamma-linoleic acid, ratio of visceral to subcutaneous adipose tissue volume, subcutaneous adipose tissue attenuation, fasting glucose, post-prandial glucose, fasting insulin, corrected insulin response, serum creatinine, diastolic blood pressure, diabetic kidney disease, type 2 diabetes and coronary artery disease- were associated nominally with signals within/near *BAI3* [24]. Owing to its pivotal role in mediating pleiotropic effects (Influencing a wide range of phenotypic traits), we strongly consider *BAI3* as a plausible player for obesity and related metabolic derangements in diverse human populations including Indians.

We also observed an association of variant rs2078267 in *SLC22A11*, a known GWAS signal for uric acid in several populations including Indians [15,52]. The SNP rs2078267 overlie an active enhancer region in subcutaneous adipose tissue and is potential binding site for numerous TFs concerned in obesity and associated metabolic disorders [34]. This SNP was also associated with differential methylation of five CpG sites within *SLC22A11* in blood. *SLC22A11* expresses in renal membranes and fetal-facing basal membrane of placenta and is a transporter for glutamate [53,54]. Higher glutamate uptake has been associated with higher BMI in Chinese adults [55]. Further, glutamate release mediates leptin action on energy expenditure in mice [56]. These findings suggest that change in expression due to genetic variations in *SLC22A11* may affect adiposity by glutamate-mediated leptin signaling.

Interestingly, we also discovered novel association of variant residing in active chromatin region - rs8100011 in *ZNF45* gene with BMI at near genome-wide significance level. ZNF45 is a predicted transcription factor [42] that binds with CEBPA, a major transcription factor for adipocyte maturation and CBX5, a histone modifier involved in proliferation and differentiation of cellular lineages including adipocytes [57,58]. The variant (rs8100011) was identified as explicit *cis*-eQTL for *ZNF45* expression in subcutaneous adipose tissue by GTEx consortium [33] and identified risk allele (A) has been associated with lower expression of gene. GTEx data is in agreement with our expression correlation analysis in Indians that show decreased expression of *ZNF45* in skeletal muscle of obese individuals and opens up further exploration into mechanistic insight of *ZNF45* variant in scheming obesity features. Although the effect of rs8100011 on obesity by modulating *ZNF45* expression seems a more plausible explanation, alternative modes of action of rs8100011 in obesity biology cannot be negated as rs8100011 was also identified as eQTL for other genes such as *ZNF155*, *ZNF283*, *ZNF404*, *AC084219.4*, *KCNN4*, *RP11-15A1.3*.

Further, imputed *ZNF45* variant - rs55736013 constituted strong binding site for several members of Krüppel-Like adipogenesis related TFs such as KLF4, KLF5 and KLF9, that promote adipogenesis via stimulating CCAAT-enhancer-binding proteins (C/EBPs) [35, 59]. Additionally, another imputed variant - rs11880216 constitute binding site for IRF3, a major transcriptional regulator of adipose inflammation, involved in maintaining systemic glucose and energy homeostasis [35,60].

Interestingly, *BAI3*, *SLC22A11* and *ZNF45* signals actively regulate multiple adiposity related traits in Indians that include - BMI, weight, WHR, WC, HC, LDL-C, TC, uric acid, and systolic blood pressure, indicating their firm control on extensive metabolic outcomes in Indians. In addition, we also found convincing evidence that discovered *BAI3*, *SLC22A11* and *ZNF45* variants displayed strong associations with numerous metabolic phenotypes in different populations [24]. These traits include BMI, weight, WC, HC, WHR, body fat percentage, fasting glucose, fasting insulin, post-prandial glucose, total cholesterol, LDL cholesterol, HDL cholesterol, leptin, adiponectin, oleic acid, creatinine, type 2 diabetes and all diabetic kidney disease. This essentially pinpoints towards their dynamic metabolic influence in diverse human populations as well.

Further, our gene-based analysis revealed additional new loci *CPS1* (Carbamoyl-Phosphate Synthase 1) and *UPP2* (Uridine Phosphorylase 2), both residing within chromosome 2 of the human genome that powerfully dictate BMI levels in Indians, and have never been accounted for obesity in earlier GWAS populations.

In conclusion, we identified three novel loci in - *BAI3*, *SLC22A11*, and *ZNF45* regulating BMI in Indians. The identification of novel genes entails for population-specific genetic studies in diverse human populations. Discovered genetic leads exclusively opens up a new biology and therapeutic considerations for obesity phenotype in genetically diverse Indians.

## Acknowledgements

The authors are greatly thankful to all the study participants. We acknowledge the support and participation of members of the INDICO consortium in data generation. We thank Prof. Mark I McCarthy for providing data for South Asian subjects of United Kingdom Biobank (UKBB) and Prof. Pawan Dhar (School of Biotechnology, Jawaharlal Nehru University, India) for critical evaluation of the manuscript. GP and AKG acknowledge University Grants Commission (UGC), Government of India for Senior Research Fellowship. KB acknowledges Council of Scientific and Industrial Research (CSIR), Government of India for Senior Research Fellowship.

## Members of INDICO

Viswanathan Mohan, Saurabh Ghosh, Abhay Sharma, Rubina Tabassum, Ganesh Chauhan, Anubha Mahajan, Om Prakash Dwivedi, Lakshmi Ramakrishnan, Radha Venkatesan, M. Chidambaram, D. Prabhakaran, K.S Reddy, Monisha Banerjee, Madhukar Saxena, Sandeep Mathur, Anil Bhansali, Viral Shah, S.V. Madhu, R.K. Marwah, Pradeep Venkatesh, S.K. Aggarwal, Shantanu SenGupta, Sreenivas Chavali, Amitabh Sharma, Analabha Basu, Khushdeep Bandesh, Anil K Giri, Shraddha Chakraborty, Yasmeen Kauser, B. Abitha, Aditya Undru, Donaka Rajashekar, Vaisak Parekatt, Suki Roy, Anjali Singh, Priyanka Banerjee, Gauri Prasad, Punam Jha, Nikhil Tandon and Dwaipayan Bharadwaj (Coordinator).

## Author Contributions

AKG and GP assembled and analyzed data; contributed to discussions and wrote the manuscript. KB provided intellectual inputs. AKG, GP, KB, VP, PB, YK, SC, DR, and INDICO helped in sample preparation and data generation. AM and AB helped in statistical analysis. AM, AS, SKM, AB, MIM and NT critically reviewed the manuscript. DB is the guarantor of this work who conceived, supervised, obtained financial support, and oversaw the entire study.

