## Supplementary Material for "Multifaceted genome-wide study identifies novel regulatory loci for body mass index in Indians"

This file includes ten tables and fifteen figures

Contents include:

**Table S1 Phenotypic characteristics of adult study subjects**

**Table S2 Association status of earlier known GWAS signals for BMI that were genotyped in replication phase in Indian adults**

**Table S3 Earlier reported associations of novel variants within *BAI3, SLC22A11* and *ZNF45***

**Table S4 In-silico replication and meta-analysis of Indian subjects with South Asian samples of UKBB cohorts**

**Table S5 eQTL analysis for *ZNF45* variant rs8100011 in GTEx browser**

**Table S6 Imputation analysis of novel loci**

**Table S7 Association of *BAI3, SLC22A11* and *ZNF45* with key metabolic traits in Indians**

**Table S8 Gene regulatory features for associated CpG sites in K562 cell line**

**Table S9 Gene based association analysis for BMI in Indians**

**Table S10 Multivariate gene-based association analysis for *BAI3* and *ZNF45* loci with adiposity measures in Indians**

**Fig.S1 Data analysis pipeline: Integration of multi-layered datasets**

**Fig.S2 Imputation analysis workflow**

**Fig.S3 Statistical power of study**

**Fig.S4 Quantile-Quantile plot (QQ-Plot) between observed and theoretical distribution of p-values in discovery phase**

**Fig.S5 Polygenic risk score analysis based on weighted and unweighted number of “effective” risk alleles**

**Fig.S6 Previous associations within/near *BAI3, SLC22A11* and *ZNF45* genes with BMI**

**Fig.S7 *ZNF45* variant rs8100011 as robust cis-eQTL in various human tissues**

**Fig.S8 Correlation among adiposity traits in Indians**

**Fig.S9 Pleiotropic effects of identified SNPs on various adiposity and metabolic features**

**Fig.S10 Gene expression profiles of *BAI3, SLC22A11* and *ZNF45* in various human tissues**

**Fig.S11 Gene regulatory features of *BAI3* locus**

**Fig.S12 Gene regulatory features of *SLC22A11* locus**

**Fig.S13 Gene regulatory features of *ZNF45* locus**

**Fig.S14 Gene expression profiles of *BAI3, SLC22A11* and *ZNF45* in human blood**

**Fig.S15 DNA methylation marks around *BAI3, SLC22A11* and *ZNF45* loci in CD14 (positive) monocytes, B-cells and T-cells**

**Table S1** Phenotypic characteristics of adult study subjects

|  | **Discovery phase** | | **Replication phase** | |
| --- | --- | --- | --- | --- |
|  | **Male (N= 597)** | **Female (N=545)** | **Male (N=2 641)** | **Female (N=2 190)** |
| **Trait** | **Median (IQR)** | **Median (IQR)** | **Median (IQR)** | **Median (IQR)** |
| Age(years) | 50 (45 - 60) | 50 (43 - 60) | 50 (43 - 60) | 48 (51-58) |
| Weight(kg) | 65.4 (54.89 - 73.95) | 60.11 (50.36 - 68.70) | 69.63 (60.11 - 79.41) | 63.94 (54.70 - 72.10) |
| BMI(kg/m2) | 23.37 (20.20 - 26.59) | 25.63 (21.79 - 29.32) | 25.19 (21.89 - 28.40) | 27.34 (23.71 - 30.80) |
| WC (cm) | 89.60 (81.55 - 96.24) | 85.47 (77.84 - 94.29) | 93.15 (85.48 - 101.36) | 87.60 (80.11 - 96.24) |
| HC(cm) | 91.94 (86.34 - 97.42) | 100.49 (91.94 - 108.68) | 97.42 (90.86 - 102.65) | 101.51 (93.07 - 108.68) |
| WHR | 0.79 (0.76 - 0.81) | 0.80 (0.78 - 0.82) | 0.94 (0.89 - 0.99) | 0.94 (0.89 - 0.98) |

Data presented as median (interquartile range-IQR) of samples

**Table S2 Association status of earlier known GWAS signals for BMI that were genotyped in replication phase in Indian adults**

|  |  |  |  |  |  |  | **Replication Phase** | | |
| --- | --- | --- | --- | --- | --- | --- | --- | --- | --- |
| **SNP** | **CHR** | **Base Position** | **Nearby Gene** | **SNP location** | **Alleles (Effect/Other)** | **MAF** | **N** | **p value** | **Effect (SE)** |
| rs17782313 | 18 | 57851097 | *MC4R* | Intergenic | G/A | 0.38 | 4569 | 3.98x10^-8^ | 0.61(0.11) |
| rs12970134 | 18 | 57884750 | *MC4R* | Intergenic | A/G | 0.37 | 4551 | 8.53 x10^-8^ | 0.59(0.11) |
| rs6265 | 11 | 27679916 | *BDNF* | missense | A/G | 0.21 | 4570 | 1.26 x10^-4^ | -0.50(0.13) |
| rs8050136 | 16 | 53816275 | *FTO* | Intronic | A/C | 0.34 | 4560 | 1.52 x10^-4^ | 0.43(0.11) |
| rs9939609 | 16 | 53820527 | *FTO* | Intronic | A/T | 0.34 | 4551 | 2.01 x10^-4^ | 0.42(0.11) |
| rs1558902 | 16 | 53803574 | *FTO* | Intronic | T/A | 0.36 | 4564 | 3.11 x10^-4^ | 0.40(0.11) |
| rs1121980 | 16 | 53809247 | *FTO* | Intronic | A/G | 0.43 | 4554 | 1.21 x10^-3^ | 0.35(0.11) |
| rs17124318 | 1 | 63480730 | *LOC199899* | Intergenic | G/C | 0.17 | 4574 | 9.78 x10^-3^ | -0.37(0.14) |
| rs10830963 | 11 | 92708710 | *MTNR1B* | Intronic | G/C | 0.41 | 4574 | 0.01 | 0.27(0.11) |
| rs7903146 | 10 | 114758349 | *TCF7L2* | Intronic | A/G | 0.31 | 4540 | 0.01 | -0.29(0.12) |
| rs174550 | 11 | 61571478 | *FADS1* | Intronic | G/A | 0.18 | 4349 | 0.01 | 0.36(0.15) |
| rs29941 | 19 | 34309532 | *KCTD15* | Intergenic | A/G | 0.39 | 4474 | 0.01 | -0.26(0.11) |
| rs7561317 | 2 | 644953 | *TMEM18* | Intergenic | A/G | 0.14 | 4555 | 0.04 | -0.32(0.16) |
| rs7498665 | 16 | 28883241 | *SH2B1* | missense | G/A | 0.24 | 4516 | 0.04 | 0.26(0.13) |

Association analysis with BMI levels, adjusted for age and sex. SNP location is in context of position with respect to the gene. MAF: Minor allele frequency. Effect size was calculated with respect to the minor allele

**Table S3 Earlier reported associations of novel variants within *BAI3, SLC22A11* and *ZNF45***

| **rs6913677 (*BAI3*)- Earlier associations** | | | |
| --- | --- | --- | --- |
| **Trait** | **Dataset** | **P value** | **Direction of effect** |
| Weight* | EXTEND GWAS | 2x10^-3^ | ↑ |
| BMI* | EXTEND GWAS | 2x10^-3^ | ↑ |
| Waist circumference* | EXTEND GWAS | 3x10^-3^ | ↑ |
| Hip circumference* | EXTEND GWAS | 3x10^-3^ | ↑ |
| Body fat percentage* | EXTEND GWAS | 0.02 | ↑ |
| Fasting glucose adj age-sex | BioMe AMP T2D GWAS | 0.02 | ↓ |
| Creatinine | GoDarts Illumina HumanOMNIExpress GWAS | 0.02 | ↑ |
| eGFR-creat (serum creatinine) | BioMe AMP T2D GWAS | 0.02 | ↓ |
| Waist-hip ratio* | EXTEND GWAS | 0.03 | ↑ |
| Fasting insulin adj BMI | Diabetic Cohort-Singapore Prospective Study GWAS | 0.04 | ↑ |
| Type 2 diabetes | EXTEND GWAS | 0.04 | ↑ |
| Cholesterol* | GLGC GWAS | 0.04 | ↑ |
| Leptin* | Leptin GWAS | 0.04 | ↑ |
| Oleic acid* | CHARGE Fatty Acid GWAS | 0.04 | ↓ |
| All diabetic kidney disease | SUMMIT Diabetic Kidney Disease GWAS: subjects with T1D or T2D | 0.04 | ↑ |
| **rs2078267 (*SLC22A11*)-Earlier associations** | | | |
| Serum urate | Global Urate Genetics Consortium GWAS | 8.7x10^-36^ | ↓ |
| Modified Stumvoll ISI adj age-sex-BMI | MAGIC GWAS | 9x10^-3^ | ↓ |
| Fasting insulin | GoT2D exome chip analysis | 0.01 | ↑ |
| Type 2 diabetes adj BMI | DIAGRAM 1000G GWAS | 0.01 | ↑ |
| Type 2 diabetes | DIAGRAM 1000G GWAS | 0.02 | ↑ |
| Oleic acid* | CHARGE Fatty Acid GWAS | 0.02 | ↓ |
| Disposition index | MAGIC GWAS | 0.04 | ↑ |
| Microalbuminuria adj HbA1c-BMI | JDRF Diabetic Nephropathy Collaborative Research Initiative GWAS | 0.04 | ↓ |
| HbA1c | EXTEND GWAS | 0.04 |  |
| **rs8100011(*ZNF45*)-Earlier associations** | | | |
| Leptin* | GoDarts Illumina Infinium GWAS | 2x10^-3^ | ↓ |
| LDL cholesterol* | GLGC GWAS | 6.8x10^-3^ | ↓ |
| Total cholesterol* | Oxford BioBank Axiom GWAS | 0.01 | ↓ |
| Cholesterol* | GLGC GWAS | 0.01 | ↓ |
| End-stage renal disease vs. macroalbuminuria adj HbA1c-BMI | JDRF Diabetic Nephropathy Collaborative Research Initiative GWAS | 0.02 | ↓ |
| Macroalbuminuria vs. controls | JDRF Diabetic Nephropathy Collaborative Research Initiative GWAS | 0.03 | ↑ |
| Two-hour glucose | MAGIC GWAS | 0.03 | ↑ |
| Macroalbuminuria vs. controls adj HbA1c-BMI | JDRF Diabetic Nephropathy Collaborative Research Initiative GWAS | 0.03 | ↑ |
| Adiponectin* | ADIPOGen GWAS | 0.04 | ↓ |
| HDL cholesterol* | Oxford BioBank Axiom GWAS | 0.04 | ↓ |

Adiposity related traits have been highlighted with * symbol. Earlier reported associations of variants with various traits were retrieved from Type 2 diabetes knowledge portal

**Table S4 In-silico replication and meta-analysis of Indian subjects with South Asian samples of UKBB cohorts**

|  | **Discovery phase in adults**  **(N = 1 142)** | | **Validation phase in adults**  **(N = 4 831)** | | **Validation phase in adolescents**  **(N =1 286)** | | **South Asians of UKBB**  **(N = 2 078)** | | **Meta-analysis (Indians+UKBB)** | | |
| --- | --- | --- | --- | --- | --- | --- | --- | --- | --- | --- | --- |
| **SNP** | **BETA**  **(SE)** | **P** | **BETA**  **(SE)** | **P** | **BETA**  **(SE)** | **P** | **BETA**  **(SE)** | **P** | **BETA**  **(SE)** | **P**  **(I^2^)** | **Dir** |
| rs6913677  (G/A)  *BAI3* | 0.71  (0.19) | 1.21 x10^-4^ | 0.38  (0.11) | 3.46 x10^-4^ | 0.55  (0.20) | 5.70 x10^-3^ | -0.02  (0.03) | 0.51 | 0.04  (0.03) | 0.19  (91.2) | +++- |
| rs2078267  (A/G)  *SLC22A11* | 0.16  (0.19) | 0.40 | 0.52  (0.10) | 6.76 x10^-7^ | 0.53  (0.20) | 7.12 x10^-3^ | 0.02  (0.03) | 0.52 | 0.07  (0.03) | 0.01  (88.9) | ++++ |
| rs8100011  (A/G)  *ZNF45* | 0.64  (0.19) | 7.17 x10^-4^ | 0.40  (0.11) | 1.76 x10^-4^ | 0.40  (0.20) | 0.05 | 0.03  (0.03) | 0.35 | 0.08  (0.03) | 6.83 x10^-3^ | ++++ |

Direction was ++/-- if there was concordance between Indian and UKBB data and +-/ -+ if there is discordance. Fixed effect inverse variance meta-analysis was done using METAL. Association results after meta-analysis of Indians and UKBB cohort has been presented. SE: Standard error; I^2^ = Chi-square value for heterogeneity test; Dir: Direction for effect size of the allele

**Table S5 eQTL analysis for *ZNF45* variant rs8100011 in GTEx browser**

| **GWAS SNP** | **Alleles** | **Gene** | ***P*** | **Effect Size** | **Tissue** | **Source** |
| --- | --- | --- | --- | --- | --- | --- |
| rs8100011 | *A/G* | ***ZNF45*** | **2.6x10^-8^** | **-0.25** | **Thyroid** | **(33)** |
|  |  | ***ZNF45*** | **1.6x10^-7^** | **-0.23** | **Adipose - Subcutaneous** | **(33)** |
|  |  | ***ZNF45*** | **1.9 x10^-7^** | **-0.17** | **Skin - Sun Exposed (Lower leg)** | **(33)** |
|  |  | ***ZNF45*** | **4.4 x10^-6^** | **-0.21** | **Nerve - Tibial** | **(33)** |
|  |  | *AC084219.4* | 2.6 x10^-5^ | 0.2 | Cells - Transformed fibroblasts | (33) |
|  |  | *KCNN4* | 5.4 x10^-6^ | 0.21 | Whole Blood | (33) |
|  |  | *RP11-15A1.3* | 2.5 x10^-6^ | 0.31 | Adipose - Visceral (Omentum) | (33) |
|  |  | *RP11-15A1.3* | 1.8 x10^-5^ | 0.23 | Artery - Tibial | (33) |
|  |  | *ZNF155* | 6.8 x10^-12^ | 0.36 | Whole Blood | (33) |
|  |  | *ZNF155* | 8.7 x10^-8^ | 0.31 | Artery - Tibial | (33) |
|  |  | *ZNF155* | 4.6 x10^-7^ | 0.38 | Adipose - Visceral (Omentum) | (33) |
|  |  | *ZNF155* | 2 x10^-6^ | 0.26 | Esophagus - Mucosa | (33) |
|  |  | *ZNF283* | 2.3 x10^-6^ | 0.33 | Brain - Cerebellar Hemisphere | (33) |
|  |  | *ZNF283* | 1.1 x10^-5^ | 0.33 | Brain - Cerebellum | (33) |
|  |  | *ZNF404* | 6.2 x10^-11^ | 0.26 | Muscle - Skeletal | (33) |
|  |  | *ZNF404* | 7.2 x10^-9^ | 0.23 | Nerve - Tibial | (33) |
|  |  | *ZNF404* | 4.5 x10^-8^ | 0.22 | Esophagus - Muscularis | (33) |
|  |  | *ZNF404* | 1.1 x10^-7^ | 0.21 | Thyroid | (33) |
|  |  | *ZNF404* | 5.7x10^-7^ | 0.2 | Adipose - Visceral (Omentum) | (33) |
|  |  | *ZNF404* | 7.2 x10^-6^ | 0.19 | Heart - Left Ventricle | (33) |
|  |  | *ZNF404* | 2.9 x10^-5^ | 0.15 | Lung | (33) |

P-value, effect size, and tissues information have been obtained by searching for rs8100011 in GTEx browser. Effect size has been represented as Z-score

**Table S6 Imputation analysis of novel loci**

***a) BAI3* signals with p-value less than or equal to genotyped signal after imputation**

| **CHR** | **SNP** | **BP** | **BETA** | **SE** | **L95** | **U95** | **P** | **Gene** | **Location** |
| --- | --- | --- | --- | --- | --- | --- | --- | --- | --- |
| 6 | rs17402611 | 69787970 | 0.7287 | 0.1848 | 0.3664 | 1.091 | 8.57x10^-5^ | *BAI3* | Intronic |
| 6 | rs9354814 | 69788137 | 0.7287 | 0.1848 | 0.3664 | 1.091 | 8.57 x10^-5^ | *BAI3* | Intronic |
| 6 | rs1336632 | 69823956 | 0.7346 | 0.1869 | 0.3683 | 1.101 | 9 x10^-5^ | *BAI3* | Intronic |
| 6 | rs4706854 | 69801042 | 0.7262 | 0.185 | 0.3637 | 1.089 | 9.15 x10^-5^ | *BAI3* | Intronic |
| 6 | rs11752858 | 69789598 | 0.7227 | 0.1846 | 0.3609 | 1.085 | 9.59 x10^-5^ | *BAI3* | Intronic |
| 6 | **rs10945151** | 69795962 | 0.718 | 0.1846 | 0.3562 | 1.08 | 1.07 x10^-4^ | *BAI3* | Intronic |
| 6 | rs1482324 | 69797221 | 0.7129 | 0.1846 | 0.3512 | 1.075 | 1.18 x10^-4^ | *BAI3* | Intronic |
| 6 | rs6913677* | 69803462 | 0.712 | 0.1846 | 0.3502 | 1.074 | 1.21 x10^-4^ | *BAI3* | Intronic |

*Index SNP. Bold imputed variants have been described in text

**b)** ***ZNF45* signals with p-value less than or equal to genotyped signal after imputation**

| **CHR** | **SNP** | **BP** | **BETA** | **SE** | **L95** | **U95** | **P** | **Gene** | **Location** |
| --- | --- | --- | --- | --- | --- | --- | --- | --- | --- |
| 19 | rs11882181 | 44437979 | -0.6628 | 0.1888 | -1.033 | -0.293 | 4.65 x10^-4^ | *ZNF45* | Intronic |
| 19 | **rs55736013** | 44437476 | -0.6552 | 0.1883 | -1.024 | -0.286 | 5.21 x10^-4^ | *ZNF45* | Intronic |
| 19 | rs55810332 | 44441752 | -0.6515 | 0.1895 | -1.023 | -0.28 | 6.09 x10^-4^ | *ZNF45* | Intergenic |
| 19 | rs8100682 | 44439937 | -0.6495 | 0.1893 | -1.021 | -0.278 | 6.24 x10^-4^ | *ZNF45* | Intergenic |
| 19 | rs10853770 | 44446951 | -0.6531 | 0.1904 | -1.026 | -0.28 | 6.27 x10^-4^ | *ZNF45* | Intergenic |
| 19 | **rs11880216** | 44444416 | -0.6502 | 0.19 | -1.023 | -0.278 | 6.45 x10^-4^ | *ZNF45* | Intergenic |
| 19 | rs8100011* | 44436733 | -0.6382 | 0.1881 | -1.007 | -0.269 | 7.17 x10^-4^ | *ZNF45* | Intronic |

*Index SNP. Bold imputed variants have been described in text

**Table S7 Association of *BAI3, SLC22A11* and *ZNF45* with key metabolic traits in Indians**

|  | | | | **Meta-analysis** | |
| --- | --- | --- | --- | --- | --- |
| **Trait** | **SNP** | **Nearby Gene** | **Alleles (Effect/Other)** | **p value** | **Effect (SE)** |
| **WHR** | rs6913677 | *BAI3* | G/A | 0.06 | 0.002 (0.001) |
|  | rs2078267 | *SLC22A11* | A/G | 0.08 | 0.002 (0.001) |
|  | rs8100011 | *ZNF45* | A/G | **5.04 × 10^−3^** | 0.004 (0.001) |
| **WC** | rs6913677 | *BAI3* | G/A | **8.75 × 10^−7^** | 1.06 (0.22) |
|  | rs2078267 | *SLC22A11* | A/G | **1.23 × 10^−5^** | 0.94 (0.22) |
|  | rs8100011 | *ZNF45* | A/G | **6.78 × 10^−8^** | 1.16 (0.22) |
| **HC** | rs6913677 | *BAI3* | G/A | **4.90 × 10^−7^** | 0.95 (0.19) |
|  | rs2078267 | *SLC22A11* | A/G | **2.30 × 10^−6^** | 0.90 (0.19) |
|  | rs8100011 | *ZNF45* | A/G | **1.20 × 10^−6^** | 0.93 (0.19) |
| **Weight** | rs6913677 | *BAI3* | G/A | **2.60 × 10^−7^** | 1.28 (0.25) |
|  | rs2078267 | *SLC22A11* | A/G | **2.25 × 10^−5^** | 1.06 (0.25) |
|  | rs8100011 | *ZNF45* | A/G | **3.70 × 10^−9^** | 1.47 (0.25) |
| **TG** | rs6913677 | *BAI3* | G/A | 0.84 | 0.26 (1.39) |
|  | rs2078267 | *SLC22A11* | A/G | 0.23 | -1.6 (1.37) |
|  | rs8100011 | *ZNF45* | A/G | 0.19 | 1.78 (1.38) |
| **LDL-C** | rs6913677 | *BAI3* | G/A | **1.73 × 10^−4^** | 2.42 (0.65) |
|  | rs2078267 | *SLC22A11* | A/G | 0.8 | -0.15 (0.64) |
|  | rs8100011 | *ZNF45* | A/G | 0.16 | 0.89 (0.64) |
| **TC** | rs6913677 | *BAI3* | G/A | **1.95 × 10^−3^** | 2.5 (0.81) |
|  | rs2078267 | *SLC22A11* | A/G | 0.82 | 0.18 (0.80) |
|  | rs8100011 | *ZNF45* | A/G | 0.063 | 1.49 (0.81) |
| **HDL-C** | rs6913677 | *BAI3* | G/A | 0.11 | 0.36 (0.23) |
|  | rs2078267 | *SLC22A11* | A/G | 0.73 | 0.08 (0.22) |
|  | rs8100011 | *ZNF45* | A/G | 0.83 | 0.05 (0.22) |
| **FG** | rs6913677 | *BAI3* | G/A | 0.12 | 0.35 (0.23) |
|  | rs2078267 | *SLC22A11* | A/G | 0.97 | -0.008 (0.22) |
|  | rs8100011 | *ZNF45* | A/G | 0.19 | 0.29 (0.22) |
| **HbA1c** | rs6913677 | *BAI3* | G/A | 0.46 | 0.009 (0.01) |
|  | rs2078267 | *SLC22A11* | A/G | 0.82 | -0.003 (0.01) |
|  | rs8100011 | *ZNF45* | A/G | 0.73 | 0.004 (0.01) |
| **FI** | rs6913677 | *BAI3* | G/A | 0.89 | 0.03 (0.18) |
|  | rs2078267 | *SLC22A11* | A/G | 0.99 | -0.002 (0.18) |
|  | rs8100011 | *ZNF45* | A/G | 0.11 | 0.28 (0.18) |
| **CRP** | rs6913677 | *BAI3* | G/A | 0.13 | -0.07 (0.05) |
|  | rs2078267 | *SLC22A11* | A/G | 0.84 | 0.009 (0.05) |
|  | rs8100011 | *ZNF45* | A/G | 0.81 | 0.01 (0.05) |
| **Urea** | rs6913677 | *BAI3* | G/A | 0.35 | 0.18 (0.20) |
|  | rs2078267 | *SLC22A11* | A/G | 0.21 | -0.24 (0.19) |
|  | rs8100011 | *ZNF45* | A/G | 0.76 | -0.06(0.20) |
| **Creatinine** | rs6913677 | *BAI3* | G/A | 0.86 | 0.008 (0.0005) |
|  | rs2078267 | *SLC22A11* | A/G | 0.74 | -0.002 (0.005) |
|  | rs8100011 | *ZNF45* | A/G | 0.07 | -0.008 (0.005) |
| **Uric acid** | rs6913677 | *BAI3* | G/A | 0.97 | 0.001(0.03) |
|  | rs2078267 | *SLC22A11* | A/G | **3.26 × 10^−11^** | -0.17(0.03) |
|  | rs8100011 | *ZNF45* | A/G | 0.62 | 0.01 (0.03) |
| **SBP** | rs6913677 | *BAI3* | G/A | **0.03** | 0.67 (0.31) |
|  | rs2078267 | *SLC22A11* | A/G | 0.51 | 0.2 (0.31) |
|  | rs8100011 | *ZNF45* | A/G | 0.91 | -0.03 (0.31) |
| **DBP** | rs6913677 | *BAI3* | G/A | 0.59 | 0.1 (0.18) |
|  | rs2078267 | *SLC22A11* | A/G | 0.2 | 0.24 (0.19) |
|  | rs8100011 | *ZNF45* | A/G | 0.42 | -0.15 (0.19) |

Association results have been obtained from meta-analysis of summary statistics of discovery and validation phases of various traits in adult cohort. Significant P-values (P < 0.05) are marked bold.

**Table S8 Gene regulatory features for associated CpG sites in K562 cell line**

| **CpG** | **Gene** | **CpG position with respect to the gene** | **Histone marks (K562)** | **K562 Chromatin state segmentation by HMM** | **Strong TF binding site** |
| --- | --- | --- | --- | --- | --- |
| cg17094144 | *BAI3* | 1stExon;5'UTR | H3K4me3, H3K27ac, H3K9ac | Active promoter | RBBP5,EZH2,MYC,RFX5,MAX |
| cg00270878 | *SLC22A11* | Gene Body, Intron | H3K4me1, H3K4me3, H3K27me3, H3K9ac | Weak enhancer | POLR2A,NFIC,KAP1,TCF7L2,TAF1,UBTF,ZNF263,GTF2F1,EP300, FOXA1,TBP,RFX5, RXRA,JUND,MYC,BHLHE40,SP1, ARID3A,MAX, RCOR1,HDAC2,FOXA2,CHD2,USF1, ATF1,SMC3,GATA3,JUN,MAZ, BRCA1,FOXP2,ATF3,CEBPD,FOS, CEBPB,FOSL2,USF2, FOXQ1 |
| cg07086353 |  | Intergenic,CTCF binding site | H3K4me1,H3K4me3, H3K27me3,H3K36me3 | Insulator | SIN3A,MYC,CTCF,TCF7L2,EZH2, YY1,GLI2,ZIC3, ZIC4,ZIC1 |
| cg08822897 |  | Intergenic | H3K27me3,H3K36me3, H3K4me1,H3K4me3,H3K9ac | Insulator | USF2,HES2,TFAP2A,TFAP2B, TFAP2C,HIC1, HIC2,KLF9 |
| cg09337943 |  | Gene Body, Intron | H3K4me1,H3K27ac, H3K27me3, H3K4me3,H3K9ac, H3K36me3 | Heterochromatin | CTCF,SP1,SP2,MZF1,TFAP2A, TFAP2B,TFAP2C |
| cg18158438 |  | TSS200 | H3K4me1,H3K4me3, H3K27ac, H3K27me3,H3K36me3, H3K9ac | Weak enhancer | CEBPB,EGR1,GLI2,Gfi1b |

The data has been retrieved for K562 cell-line from ENCODE database through UCSC genome browser

TSS200: 200 bases upstream from transcription start site. Active histone marks for gene transcription: H2K27ac, H3K4me1, H3K9ac, H3K4me3, and H3K36me3. Repressive histone mark: H3K27me3. TF: transcription factor

**Table S9 Gene based association analysis for BMI in Indians**

| **Gene** | **Gene-based assoc. P** | **Chr.** | **Gene Start_Position** | **Gene Biotype** | **Number of SNPs** | **Minimum SNP-based assoc. P** |
| --- | --- | --- | --- | --- | --- | --- |
| *CPS1* | 1.88x10^-9^ | 2 | 211458078 | protein-coding gene | 16 | 1.31x10^-6^ |
| *UPP2* | 6.80 x10^-8^ | 2 | 158958381 | protein-coding gene | 13 | 1.15x10^-4^ |
| *LINC00358* | 4.22 x10^-7^ | 13 | 62577657 | non-coding RNA | 4 | 2.99 x10^-4^ |
| *ACOXL* | 4.67 x10^-7^ | 2 | 111490149 | protein-coding gene | 29 | 1.09 x10^-4^ |
| *FAM71E2* | 1.02 x10^-6^ | 19 | 55866275 | protein-coding gene | 3 | 1.26 x10^-4^ |
| *NRCAM* | 3.11 x10^-6^ | 7 | 107788070 | protein-coding gene | 25 | 3.51 x10^-4^ |
| *LINC01142* | 5.26 x10^-6^ | 1 | 170240545 | non-coding RNA | 3 | 1.73 x10^-4^ |
| *SLC25A12* | 6.50 x10^-6^ | 2 | 172639914 | protein-coding gene | 13 | 1.29 x10^-5^ |
| *PKD1L3* | 7.55 x10^-6^ | 16 | 71963440 | protein-coding gene | 5 | 6.52 x10^-5^ |
| *SLC16A12-AS1* | 7.64 x10^-6^ | 10 | 91215747 | non-coding RNA | 10 | 1.13 x10^-3^ |
| *UBA5* | 8.24 x10^-6^ | 3 | 132379540 | protein-coding gene | 4 | 3.38 x10^-4^ |
| *APBA2* | 9.59 x10^-6^ | 15 | 29362608 | protein-coding gene | 17 | 4.36 x10^-4^ |

Variants with p < 10^-5^ have been represented. Accumulated number of markers for each gene has been shown

**Table S10 Multivariate gene-based association analysis for *BAI3* and *ZNF45* loci with adiposity measures in Indians**

| **Gene** | **Gene-based assoc. P** | **Chr** | **Gene Start_Position** | **Studied SNPs** | **SNP location** | **TASTE_P** | **SNP-based assoc. P (BMI)** | **SNP-based assoc. P (Weight)** | **SNP-based assoc. P (WHR)** | **SNP-based assoc. P (WC)** | **SNP-based assoc. P (HC)** |
| --- | --- | --- | --- | --- | --- | --- | --- | --- | --- | --- | --- |
| *BAI3* | 0.01 | 6 | 69345631 | rs6913677 | intronic | 1.38 x10^-5^ | 1.21 x10^-4^ | 2.16 x10^-4^ | 5.44 x10^-4^ | 1.38 x10^-5^ | 1.46 x10^-3^ |
|  |  |  |  | rs3799037 | intronic | 2.66 x10^-4^ | 2.66 x10^-4^ | 3.42 x10^-3^ | 6.20 x10^-3^ | 4.43 x10^-4^ | 0.02 |
|  |  |  |  | rs9363983 | intronic | 2.45 x10^-4^ | 6.36 x10^-4^ | 1.01 x10^-3^ | 2.77 x10^-3^ | 2.45 x10^-4^ | 9.12 x10^-3^ |
|  |  |  |  | rs10455681 | intronic | 8.62 x10^-5^ | 6.36 x10^-4^ | 8.33 x10^-4^ | 5.25 x10^-4^ | 8.62 x10^-5^ | 8.68 x10^-3^ |
|  |  |  |  | rs12665384 | intronic | 1.07 x10^-3^ | 1.07 x10^-3^ | 6.50 x10^-3^ | 0.04 | 3.94 x10^-3^ | 0.01 |
|  |  |  |  | rs9446089 | intronic | 4.48 x10^-4^ | 1.15 x10^-3^ | 1.49 x10^-3^ | 4.48 x10^-4^ | 4.56 x10^-4^ | 0.04 |
|  |  |  |  | rs3799019 | intronic | 2.56 x10^-3^ | 2.56 x10^-3^ | 0.02 | 0.07 | 0.01 | 0.04 |
|  |  |  |  | rs1336653 | intronic | 3.74 x10^-3^ | 3.74 x10^-3^ | 0.05 | 0.16 | 0.06 | 0.26 |
|  |  |  |  | rs1482326 | intronic | 2.69 x10^-3^ | 5.18 x10^-3^ | 0.02 | 5.36 x10^-3^ | 2.69 x10^-3^ | 0.08 |
|  |  |  |  | rs634371 | intronic | 3.73 x10^-3^ | 5.59 x10^-3^ | 0.03 | 3.73 x10^-3^ | 7.22 x10^-3^ | 0.18 |
|  |  |  |  | rs6904267 | intronic | 6.12 x10^-3^ | 6.12 x10^-3^ | 0.04 | 9.77 x10^-3^ | 0.02 | 0.21 |
|  |  |  |  | rs7756040 | intronic | 6.41 x10^-3^ | 6.41 x10^-3^ | 0.04 | 9.34 x10^-3^ | 0.02 | 0.22 |
|  |  |  |  | rs2210867 | intronic | 6.41 x10^-3^ | 6.41 x10^-3^ | 0.04 | 9.34 x10^-3^ | 0.02 | 0.22 |
|  |  |  |  | rs12213405 | intronic | 6.74 x10^-3^ | 6.74 x10^-3^ | 0.04 | 0.01 | 0.02 | 0.22 |
|  |  |  |  | rs526898 | intronic | 3.59 x10^-3^ | 9.76 x10^-3^ | 0.05 | 3.59 x10^-3^ | 0.01 | 0.25 |
|  |  |  |  | rs9454664 | intronic | 0.01 | 0.01 | 0.06 | 0.08 | 0.06 | 0.39 |
|  |  |  |  | rs3798977 | intronic | 0.01 | 0.01 | 0.08 | 0.02 | 0.03 | 0.46 |
|  |  |  |  | rs10485436 | intronic | 0.01 | 0.01 | 0.03 | 0.55 | 0.03 | 0.02 |
|  |  |  |  | rs3823070 | intronic | 0.01 | 0.01 | 0.02 | 0.05 | 0.01 | 0.04 |
|  |  |  |  | rs9454667 | intronic | 0.01 | 0.01 | 0.07 | 0.08 | 0.05 | 0.38 |
|  |  |  |  | rs7759377 | intronic | 4.17 x10^-3^ | 0.02 | 4.1x10^-3^ | 0.19 | 5.74 x10^-3^ | 4.82 x10^-3^ |
|  |  |  |  | rs12527597 | intronic | 4.17 x10^-3^ | 0.02 | 4.17x10^-3^ | 0.19 | 5.74 x10^-3^ | 4.82 x10^-3^ |
|  |  |  |  | rs7748808 | intronic | 5.52 x10^-3^ | 0.02 | 5.57 x10^-3^ | 0.21 | 7.30 x10^-3^ | 5.52 x10^-3^ |
|  |  |  |  | rs779481 | intronic | 0.02 | 0.02 | 0.07 | 0.08 | 0.05 | 0.25 |
|  |  |  |  | rs1410700 | intronic | 0.02 | 0.02 | 0.05 | 0.12 | 0.03 | 0.09 |
|  |  |  |  | rs9294819 | intronic | 0.03 | 0.03 | 0.06 | 0.07 | 0.03 | 0.11 |
|  |  |  |  | rs3799055 | intronic | 0.01 | 0.03 | 0.03 | 0.22 | 0.01 | 0.04 |
|  |  |  |  | rs17748715 | intronic | 0.03 | 0.03 | 0.06 | 0.27 | 0.08 | 0.10 |
|  |  |  |  | rs2225803 | intronic | 0.03 | 0.03 | 0.13 | 0.24 | 0.13 | 0.40 |
|  |  |  |  | rs9454693 | intronic | 0.03 | 0.03 | 0.07 | 0.04 | 0.03 | 0.15 |
|  |  |  |  | rs13191563 | intronic | 0.03 | 0.03 | 0.12 | 0.22 | 0.10 | 0.35 |
|  |  |  |  | rs1296346 | intronic | 0.04 | 0.04 | 0.07 | 0.28 | 0.09 | 0.13 |
|  |  |  |  | rs9446078 | intronic | 0.04 | 0.04 | 0.22 | 0.14 | 0.19 | 0.76 |
|  |  |  |  | rs3823061 | intronic | 0.05 | 0.05 | 0.20 | 0.84 | 0.28 | 0.14 |
|  |  |  |  | rs1952435 | intronic | 0.05 | 0.05 | 0.08 | 0.18 | 0.07 | 0.16 |
|  |  |  |  | rs3798984 | intronic | 0.02 | 0.11 | 0.02 | 0.46 | 0.07 | 0.05 |
|  |  |  |  | rs2073135 | intronic | 0.05 | 0.12 | 0.05 | 0.51 | 0.09 | 0.08 |
|  |  |  |  | rs779484 | intronic | 0.04 | 0.22 | 0.47 | 0.04 | 0.12 | 0.68 |
|  |  |  |  | rs11751520 | intronic | 0.03 | 0.25 | 0.15 | 0.62 | 0.21 | 0.03 |
|  |  |  |  | rs2785575 | intronic | 0.04 | 0.25 | 0.14 | 0.04 | 0.10 | 0.59 |
|  |  |  |  | rs6455306 | intronic | 0.05 | 0.47 | 0.83 | 0.05 | 0.35 | 0.80 |
| *ZNF45* | 5.5x10^-4^ | 19 | 44416775 | rs8100011 | intronic | 1.99 x10^-4^ | 7.17 x10^-4^ | 1.99 x10^-4^ | 0.09 | 2.17 x10^-4^ | 4.99 x10^-4^ |
|  |  |  |  | rs444816 | intronic | 2.94 x10^-5^ | 9.04 x10^-3^ | 2.94 x10^-5^ | 0.06 | 1.32 x10^-4^ | 5.65 x10^-4^ |
|  |  |  |  | rs388706 | exonic | 1.86 x10^-3^ | 0.01 | 1.86 x10^-3^ | 0.91 | 0.01 | 2.35 x10^-3^ |
|  |  |  |  | rs399098 | exonic | 6.25 x10^-4^ | 0.02 | 8.04 x10^-4^ | 0.85 | 2.98 x10^-3^ | 6.25 x10^-4^ |
|  |  |  |  | rs454376 | intronic | 2.03 x10^-3^ | 0.05 | 2.03 x10^-3^ | 0.11 | 4.10 x10^-3^ | 0.02 |

Multivariate gene-based association test by extended Simes procedure (MGAS). Association p values of markers within 2 Mb regions of *BAI3* and *ZNF45* loci for adiposity indices and trait correlation information were incorporated in MGAS based model using KGG v4

**Fig.S1 Data analysis pipeline: Integration of multi-layered datasets**

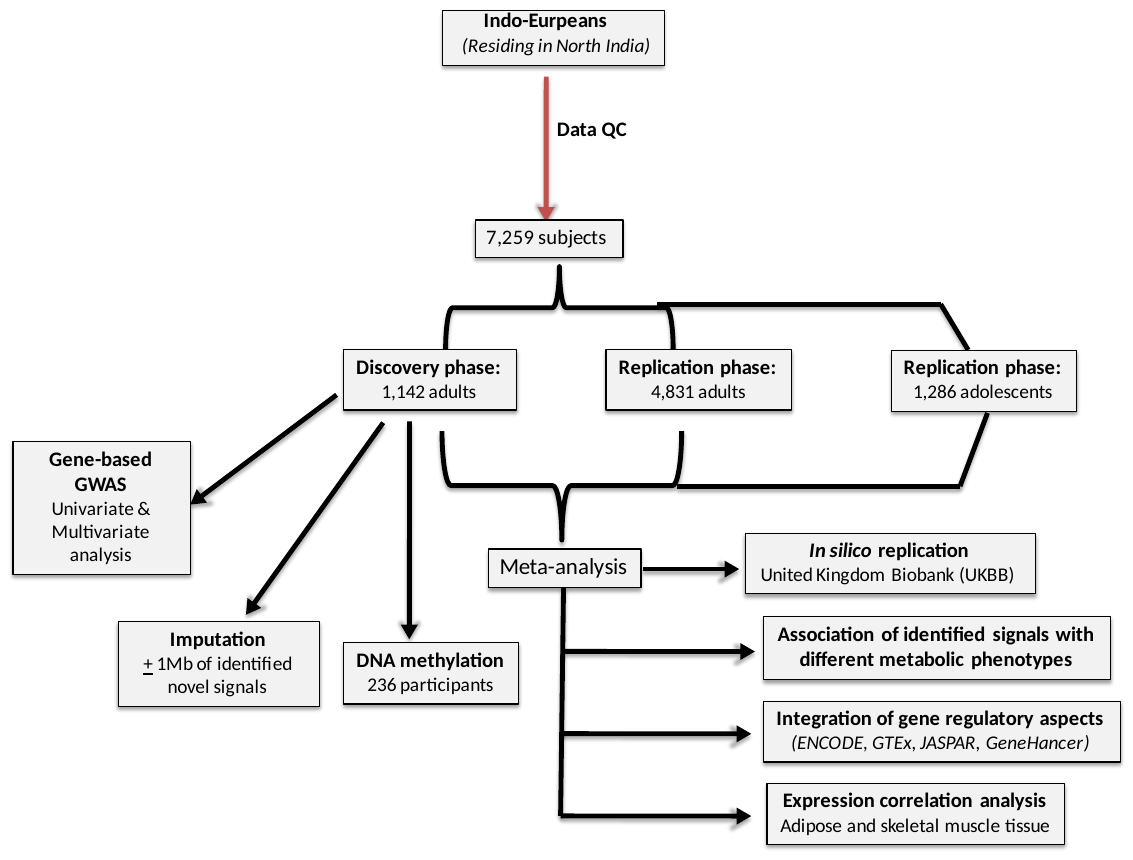

**Fig.S2 Imputation analysis workflow**

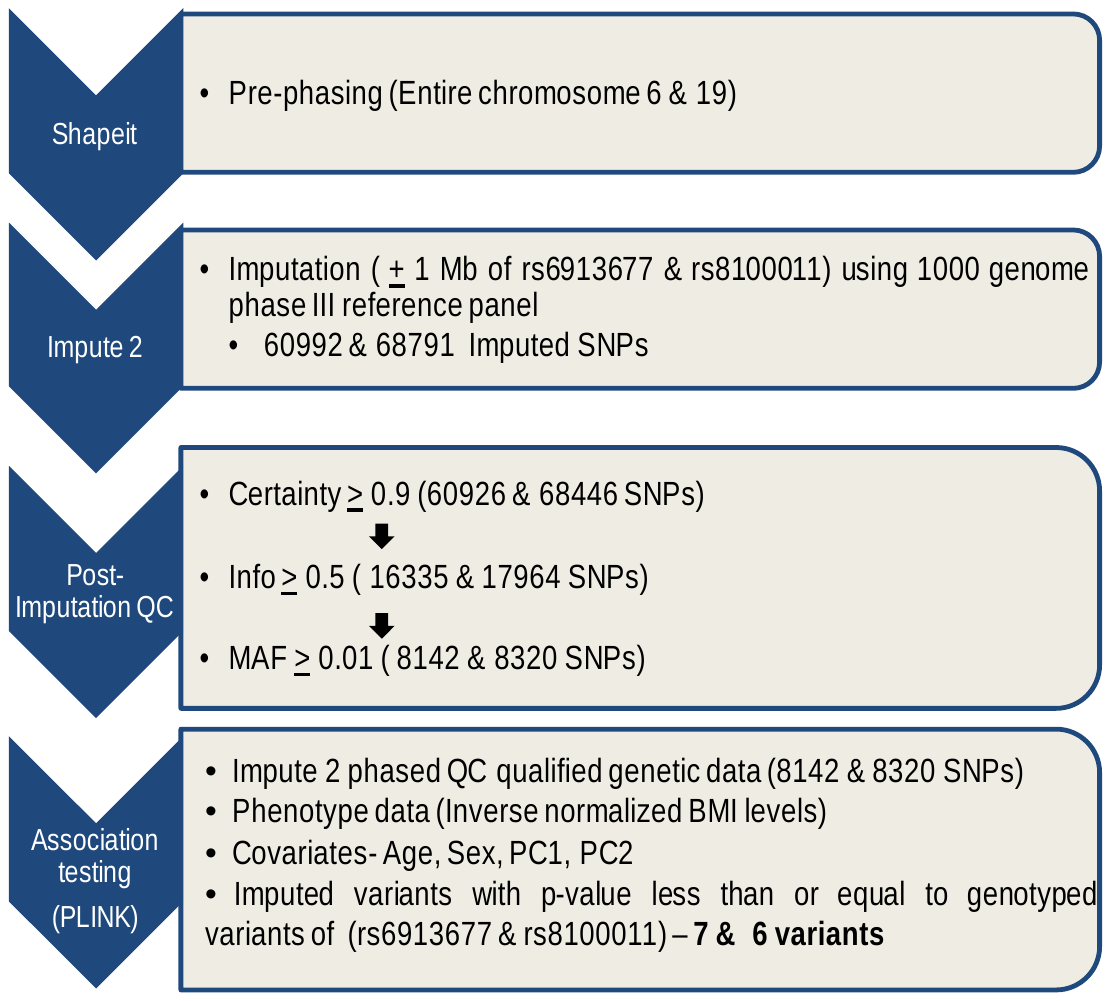

**Fig.S3 Statistical power of study**

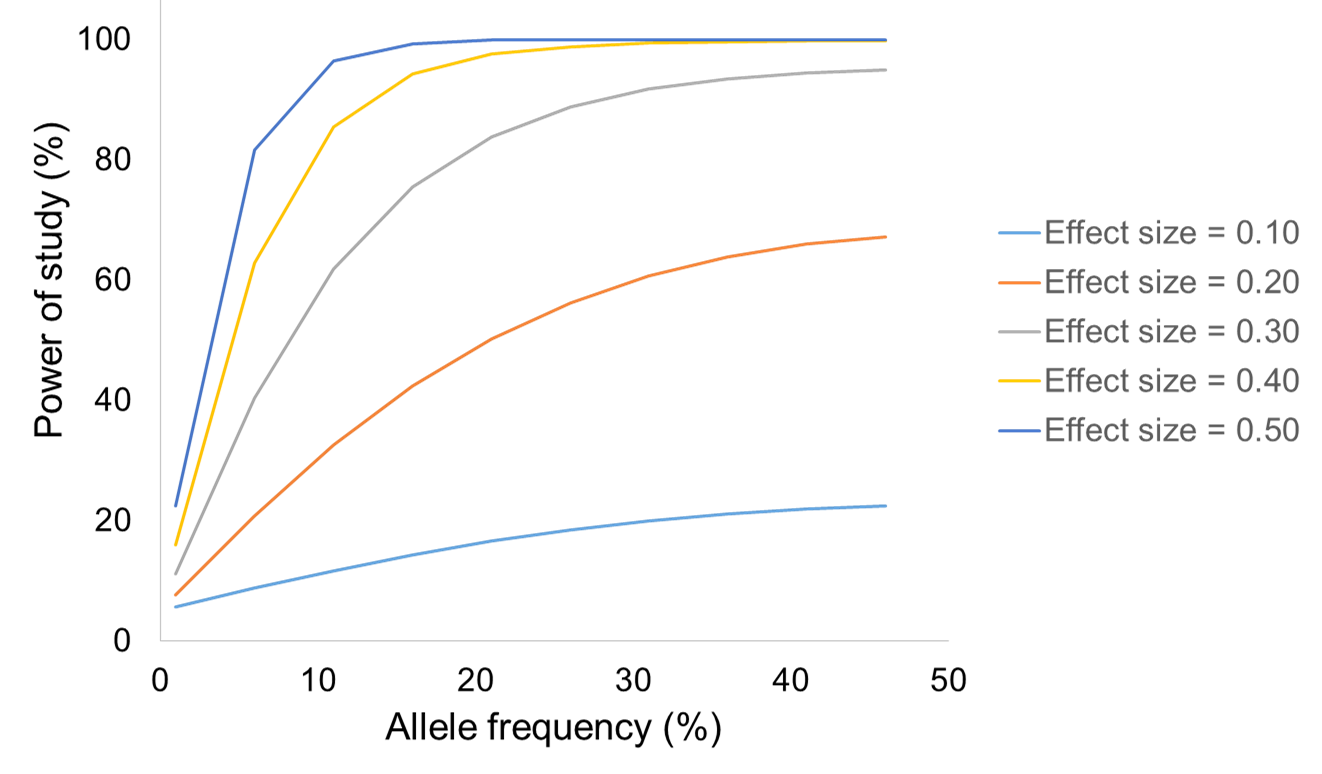

**Fig.S3** The statistical power of the study for meta-analysis was calculated for allele frequencies ranging from 0.01 to 0.50 for effect sizes from 0.1-0.5 assuming a log-additive model of inheritance at p-value = 0.05

**Fig.S4 Quantile-Quantile plot (QQ-Plot) between observed and theoretical distribution of p-values in discovery phase**

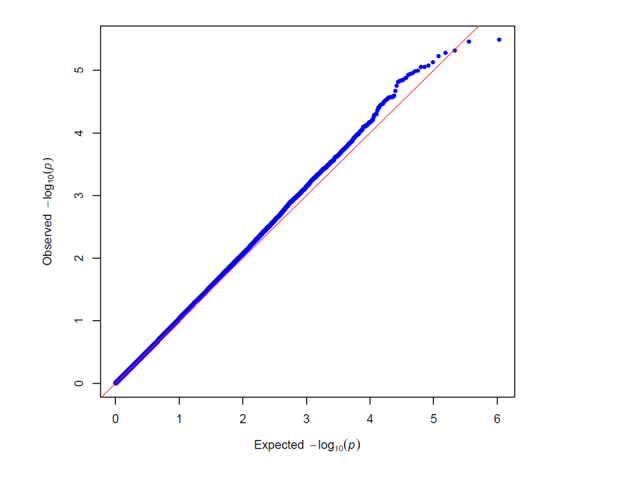

**Fig.S4** Quantile-quantile (QQ) plot for the calculated p-value in discovery phase. The -log10 of p-value observed for the association of SNPs under additive model adjusted for age, sex, PC1 and PC2 (blue symbols) are plotted against the theoretical -log10 p-value expected under the null hypothesis (red line). The genomic control inflation factor (λ) was estimated to be 1.06

**Fig.S5 Polygenic risk score analysis based on weighted and unweighted number of “effective” risk alleles**

a) **Cumulative effect of weighted number of “effective” risk alleles on BMI and risk of overweight/obesity**

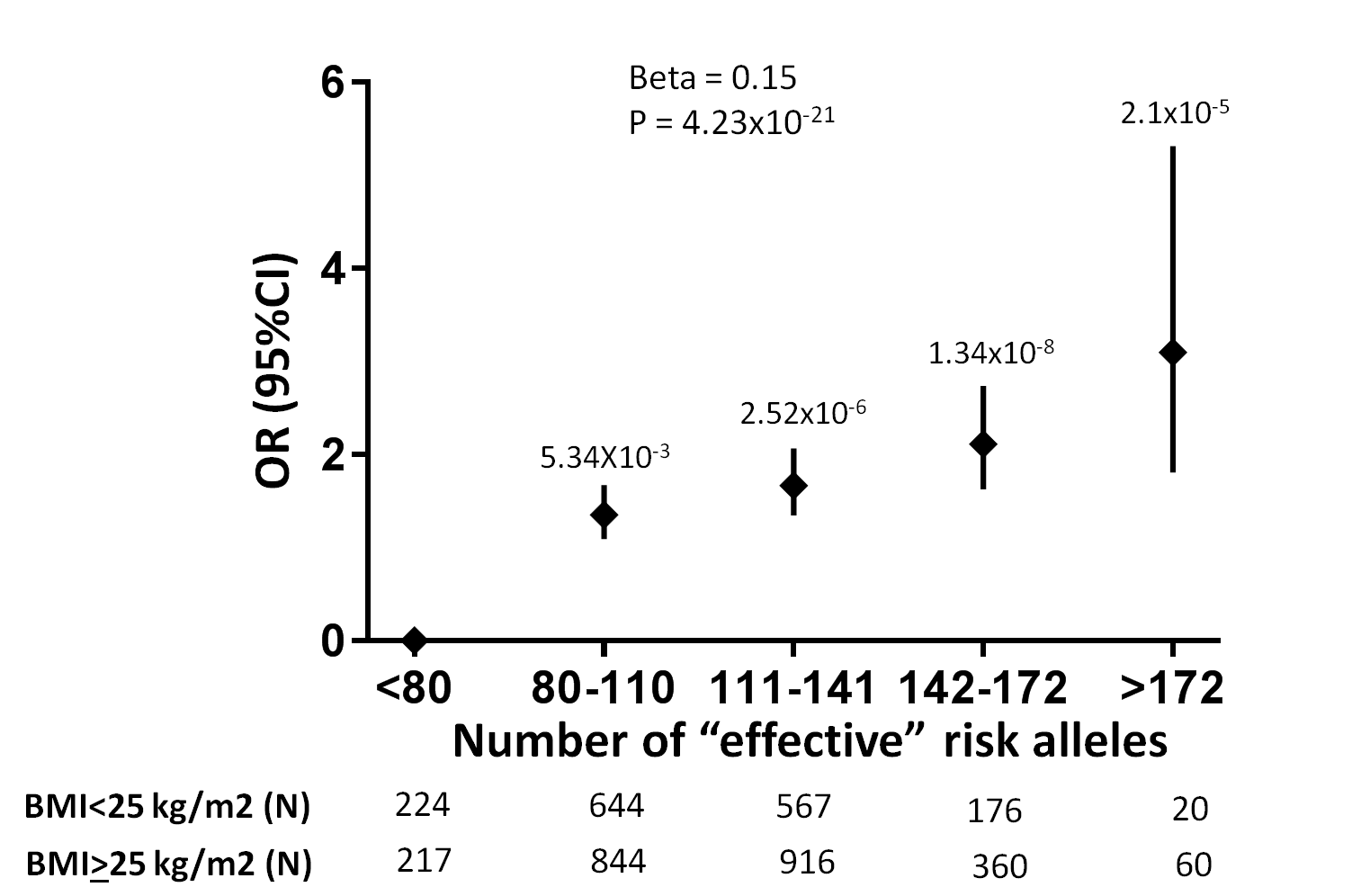

**Fig.S5 (a)** Risk alleles of 17 SNPs (14 known and 3 novel) were summed across subjects in an additive, weighted genetic model. Effect size (beta) for overall trend was computed in a linear regression model for all study subjects in which genotype data for all 17 SNPs were available. Subjects with less than 80 risk alleles have been used as a reference group to calculate risk of overweight/obesity in different risk score groups. Number of subjects with BMI< 25 and BMI≥ 25 carrying corresponding number of risk alleles has been shown below each group. Different risk allele carrying groups with two BMI strata have been compared using chi-squared test.

**b) Cumulative effect of unweighted number of “effective” risk alleles on BMI and risk of overweight/obesity**

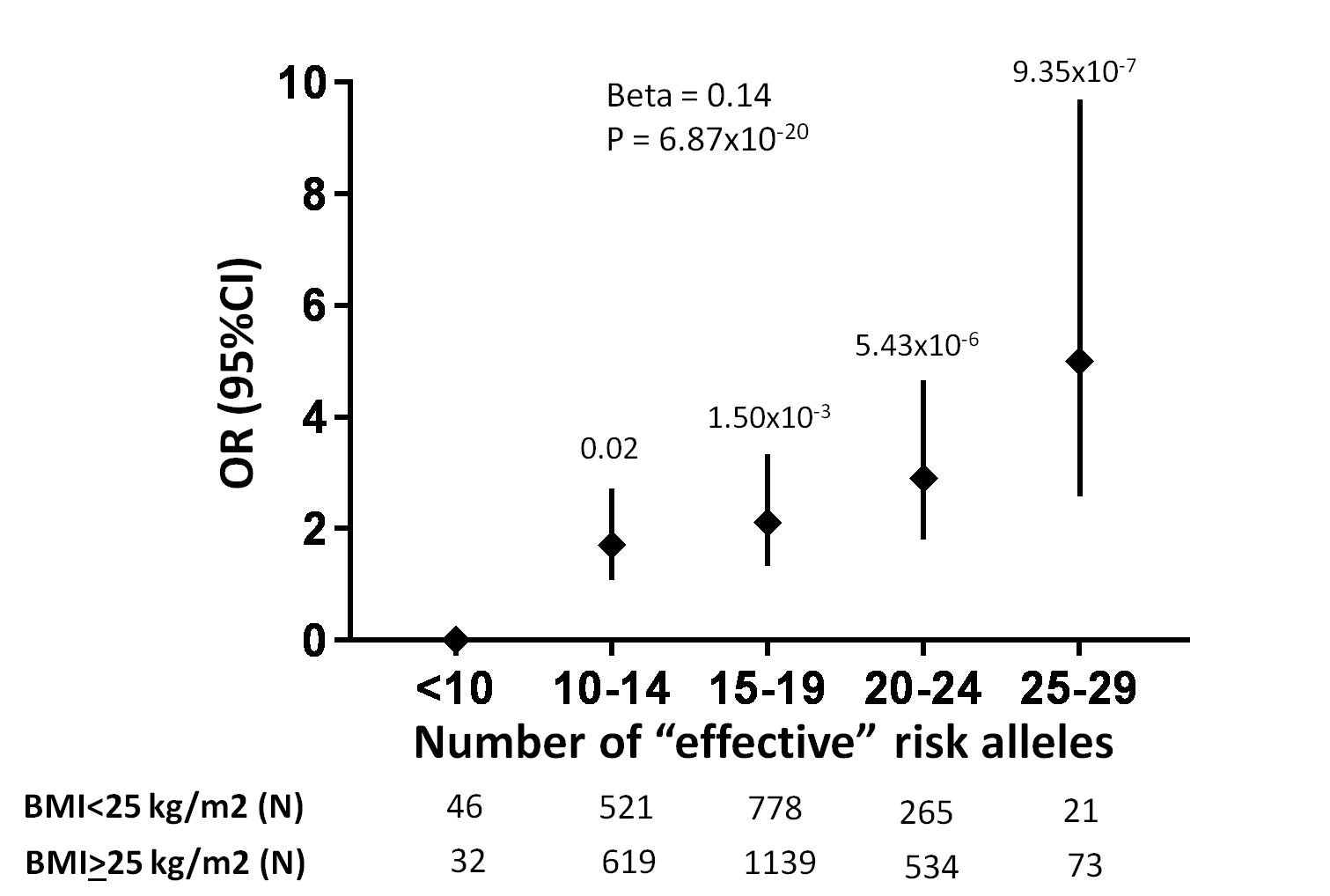

**Fig.S5 (b)** Risk alleles of 17 SNPs (14 known and 3 novel) were summed across subjects in an additive, unweighted genetic model. Effect size (beta) for overall trend was computed in a linear regression model for all study subjects in which genotype data for all 17 SNPs were available. Subjects with less than 10 risk alleles have been used as a reference group to calculate risk of overweight/obesity in different risk score groups. Number of subjects with BMI< 25 and BMI≥ 25 carrying corresponding number of risk alleles has been shown below each group. Different risk allele carrying groups with two BMI strata have been compared using chi-squared test.

**Fig.S6 Previous genetic associations within/near *BAI3, SLC22A11* and *ZNF45* genes with BMI**

a) ***BAI3***

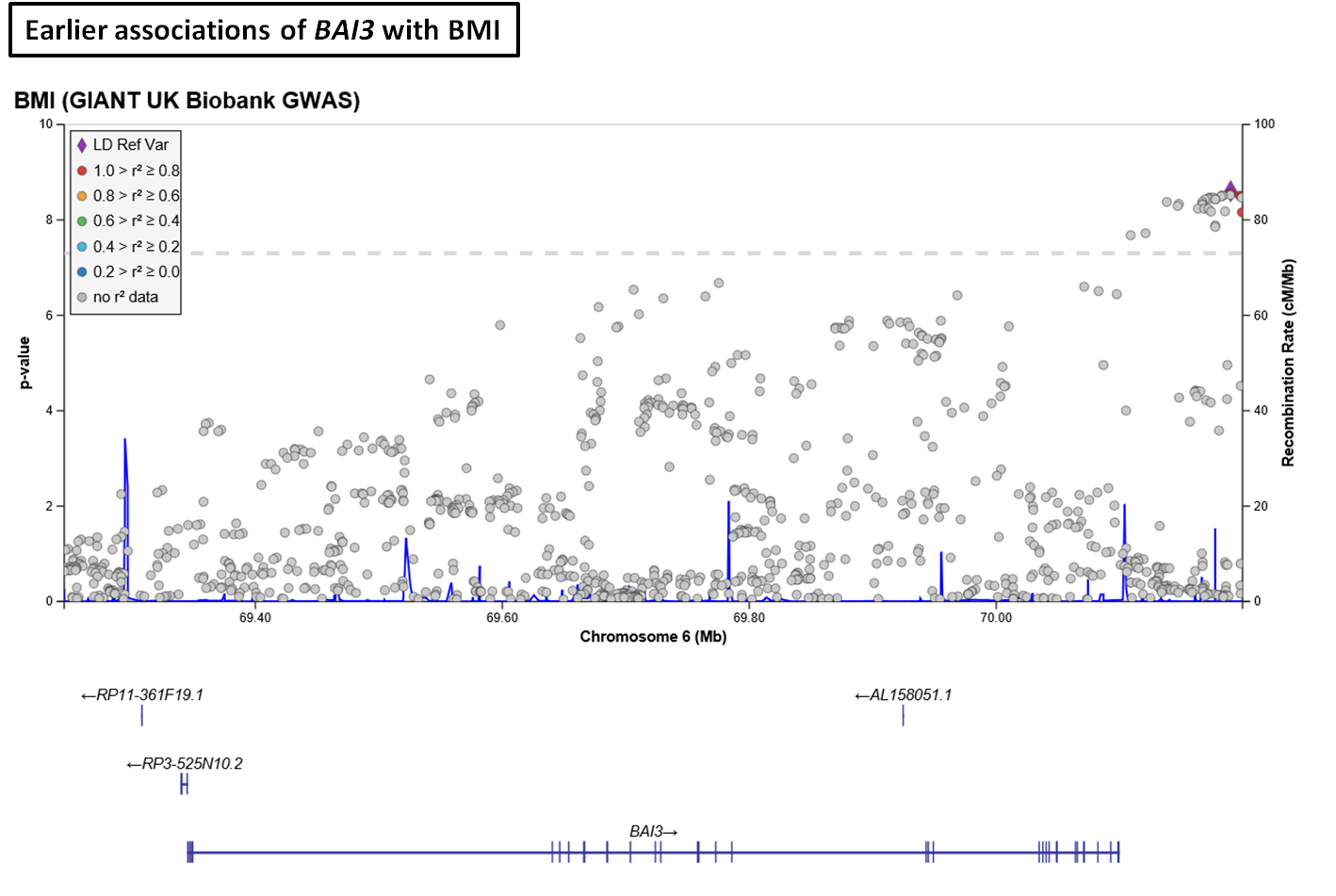

**b)** ***SLC22A11***

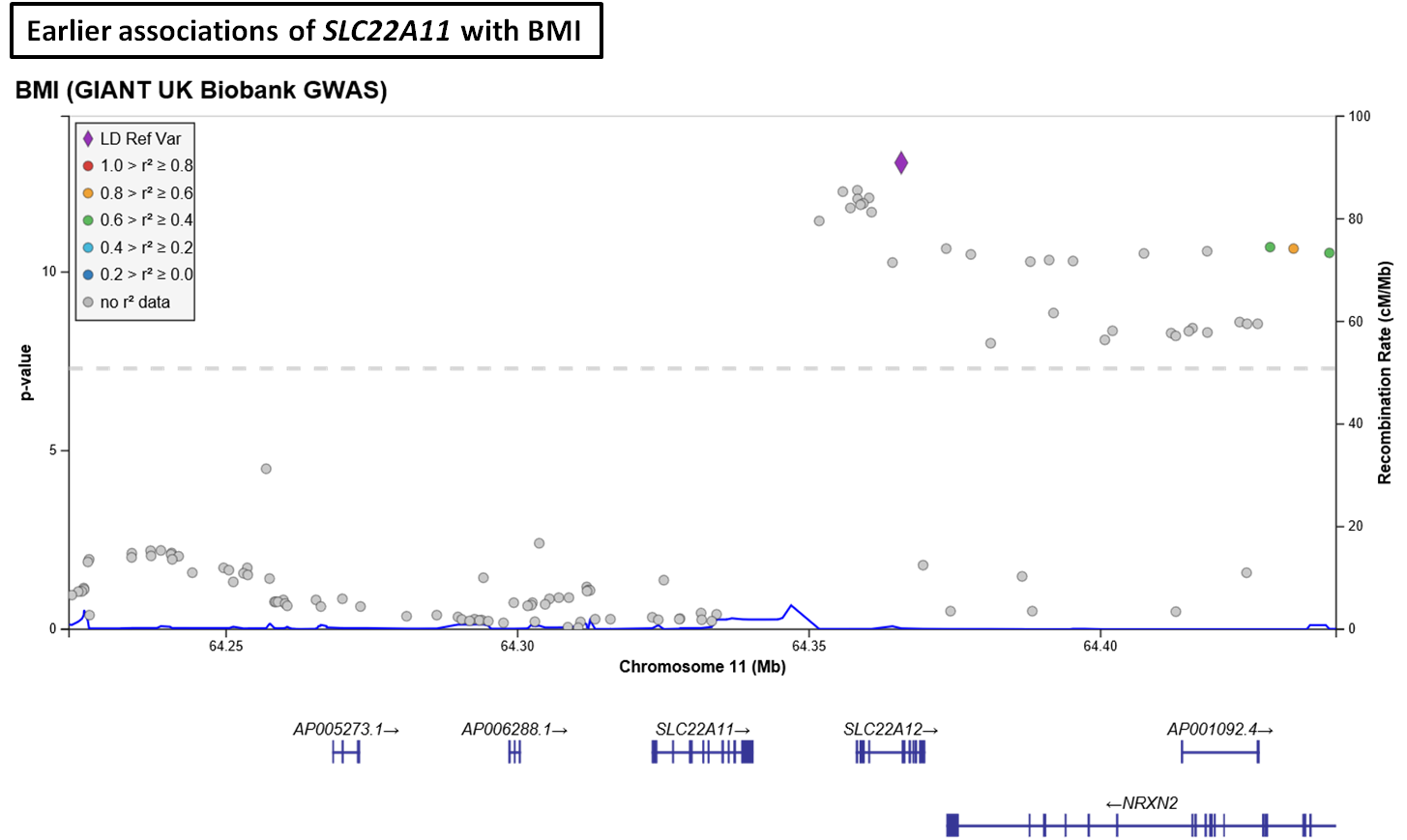

**c) *ZNF45***

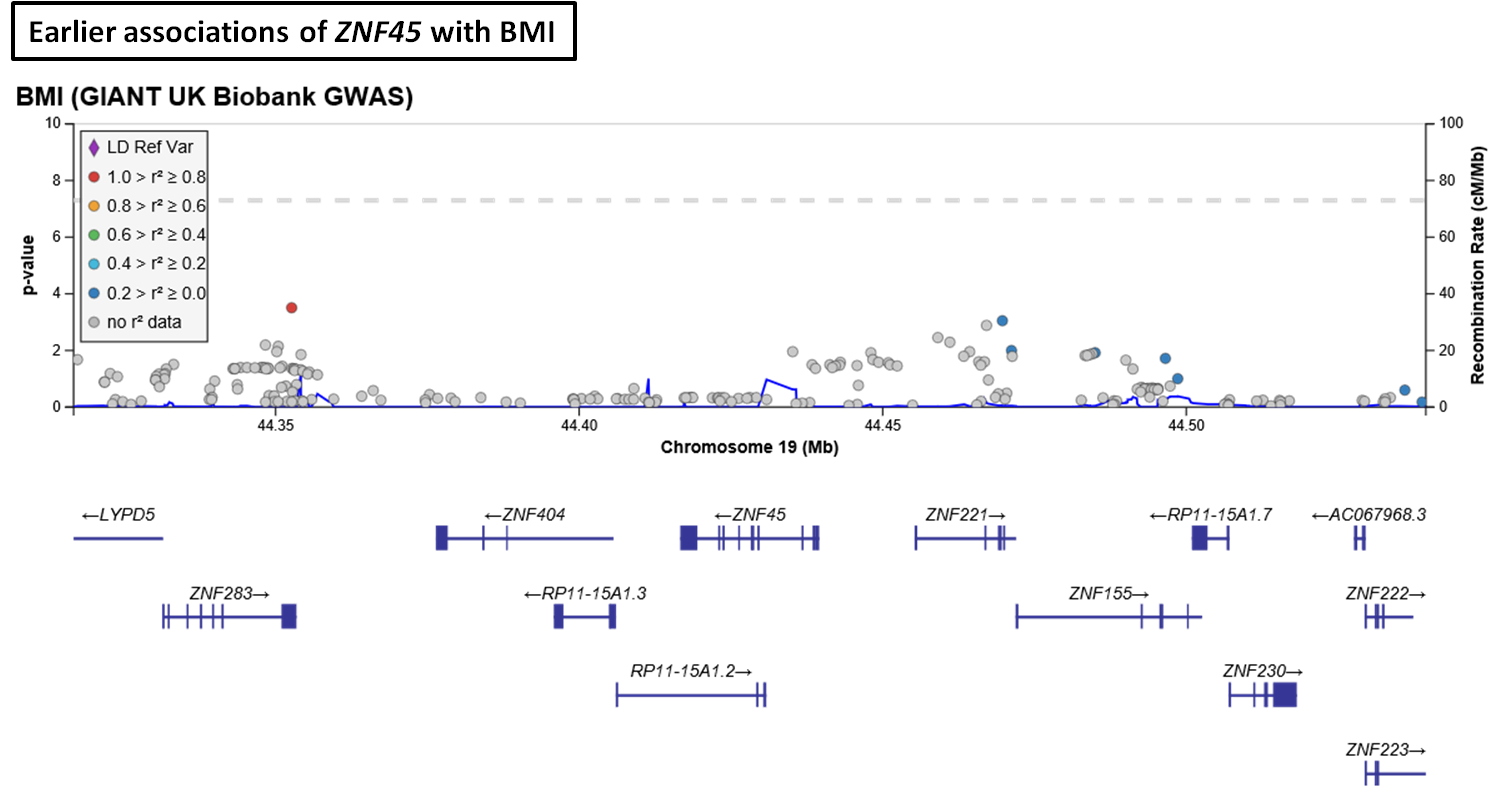

**Fig.S6** Earlier reported associations of *BAI3, SLC22A11* and *ZNF45* with BMI were derived from Type 2 diabetes knowledge portal

**Fig.S7 *ZNF45* variant rs8100011 as robust cis-eQTL in various human tissues**

**Fig.S7**

Gene expression quantitative trait loci data (eQTL data) for SNPs in human subcutaneous adipose tissue, thyroid, skin and tibial nerve were accessed from GTEx portal

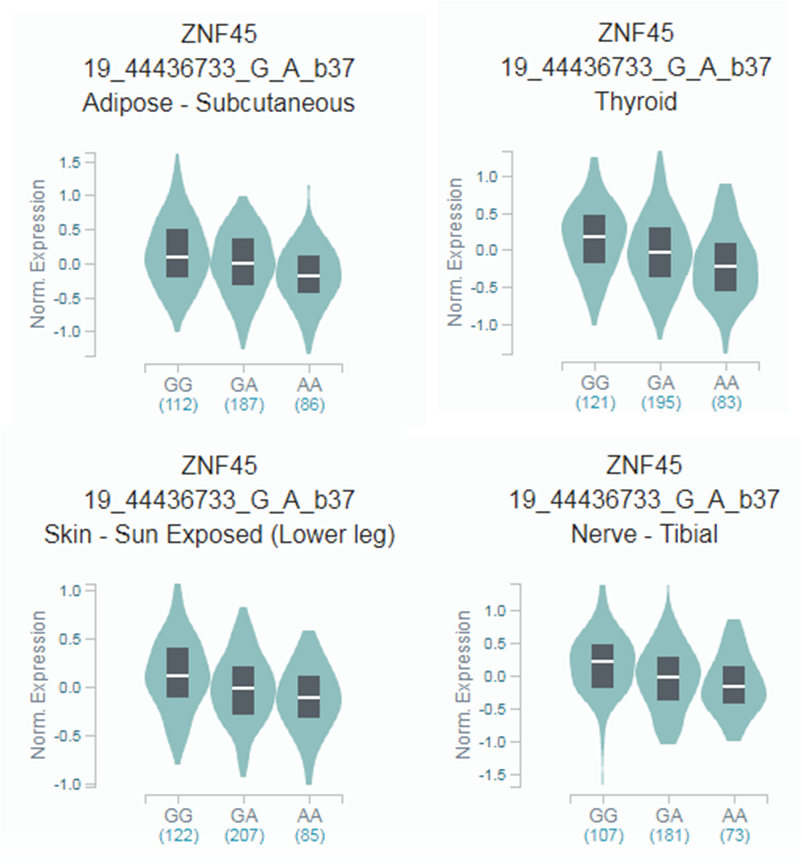

**Fig.S8 Pair wise correlation among adiposity traits in Indians**

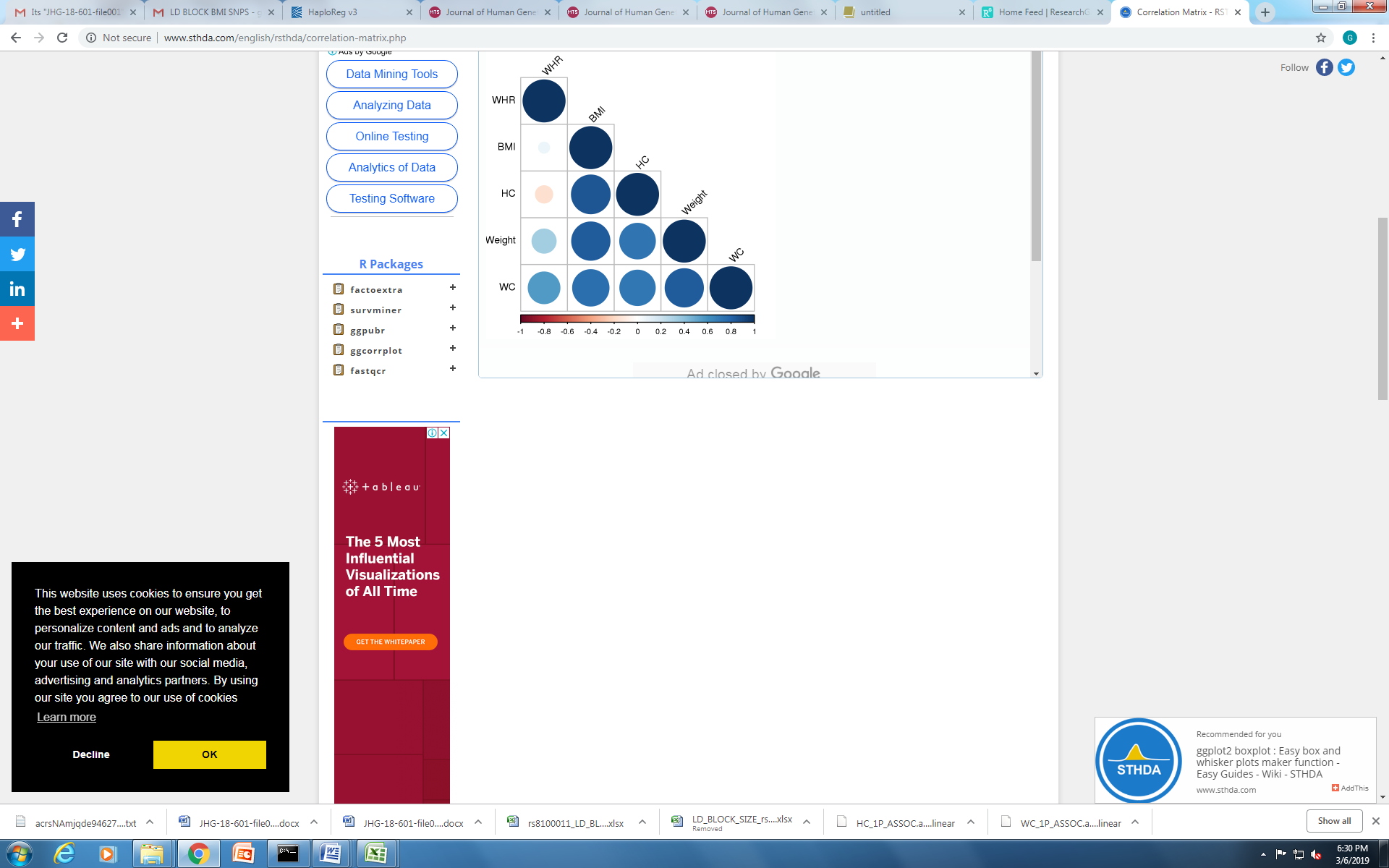

**Fig.S8**

**Figure presents pair wise correlations between adiposity traits. Positive correlations are in blue and negative correlations are in red. Color intensity and size of the circle are relative to the correlation coefficients. Color intensity scale has been shown in bottom**

**Fig.S9 Pleiotropic effects of identified SNPs on various adiposity and metabolic features**

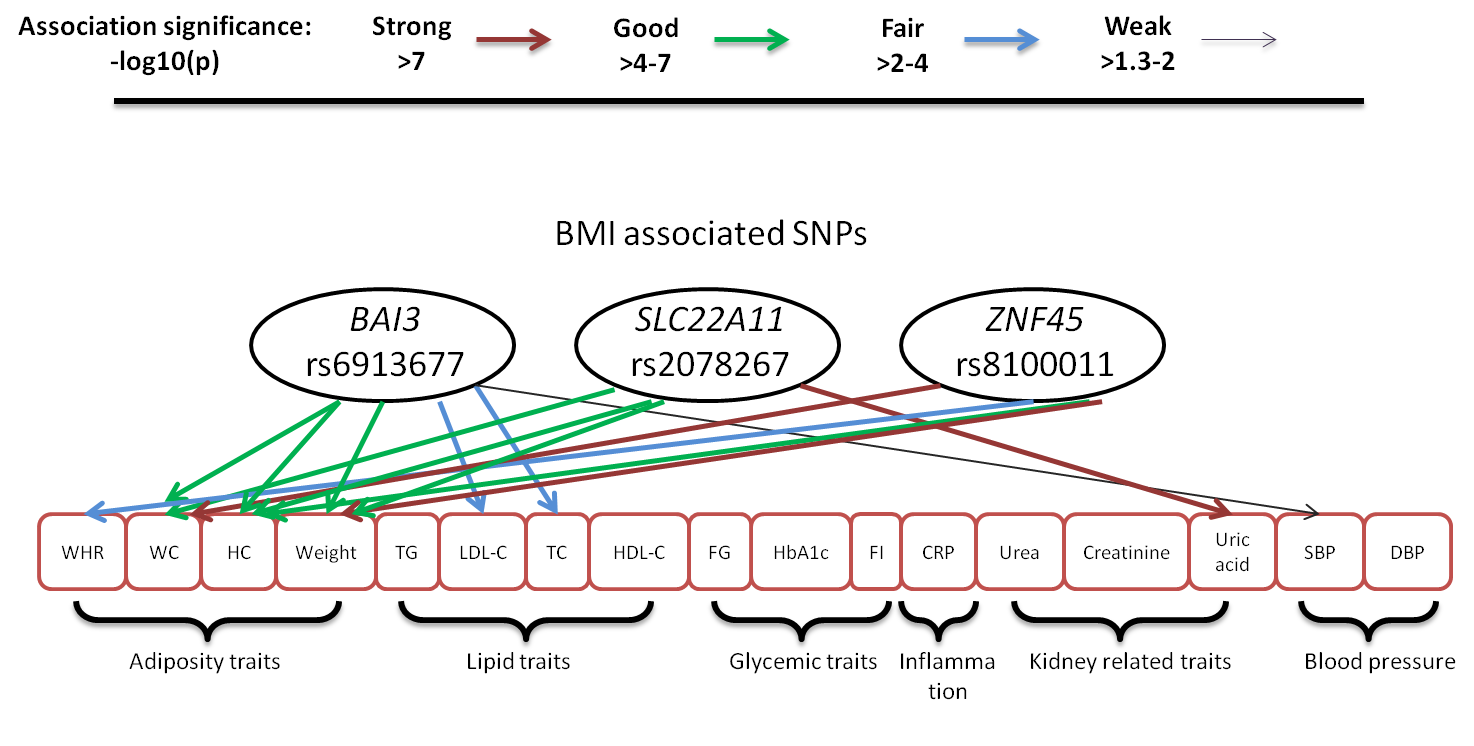

**Fig.S10 Gene expression profiles of *BAI3, SLC22A11* and *ZNF45* in various human tissues**

a) ***BAI3***

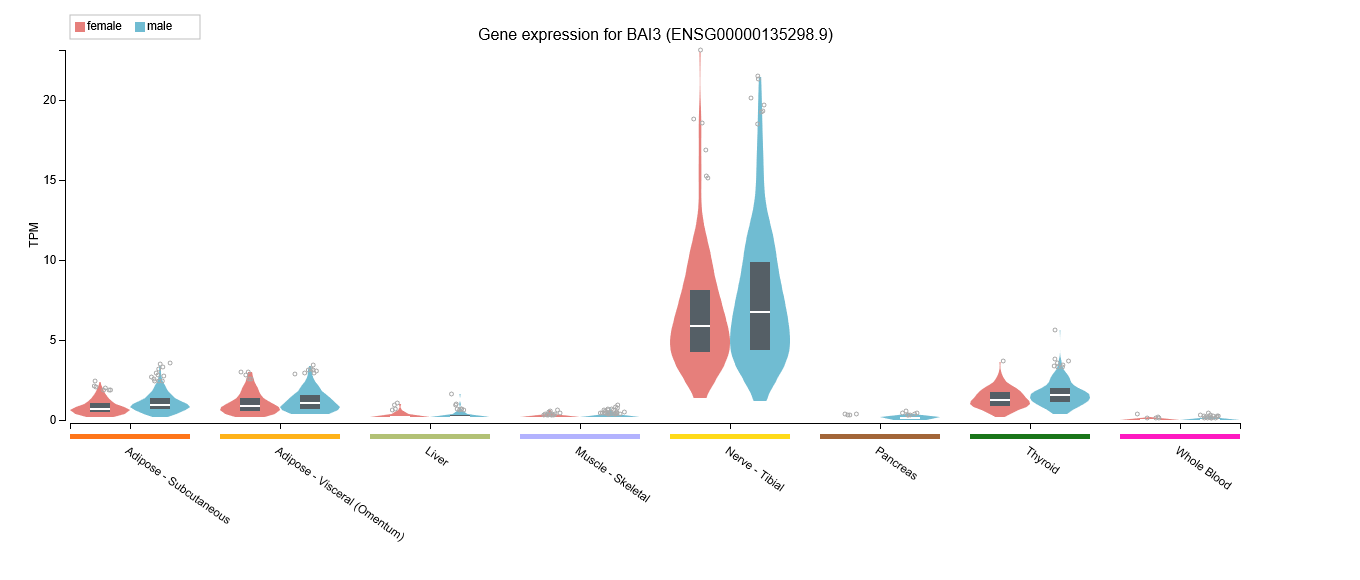

b) ***SLC22A11***

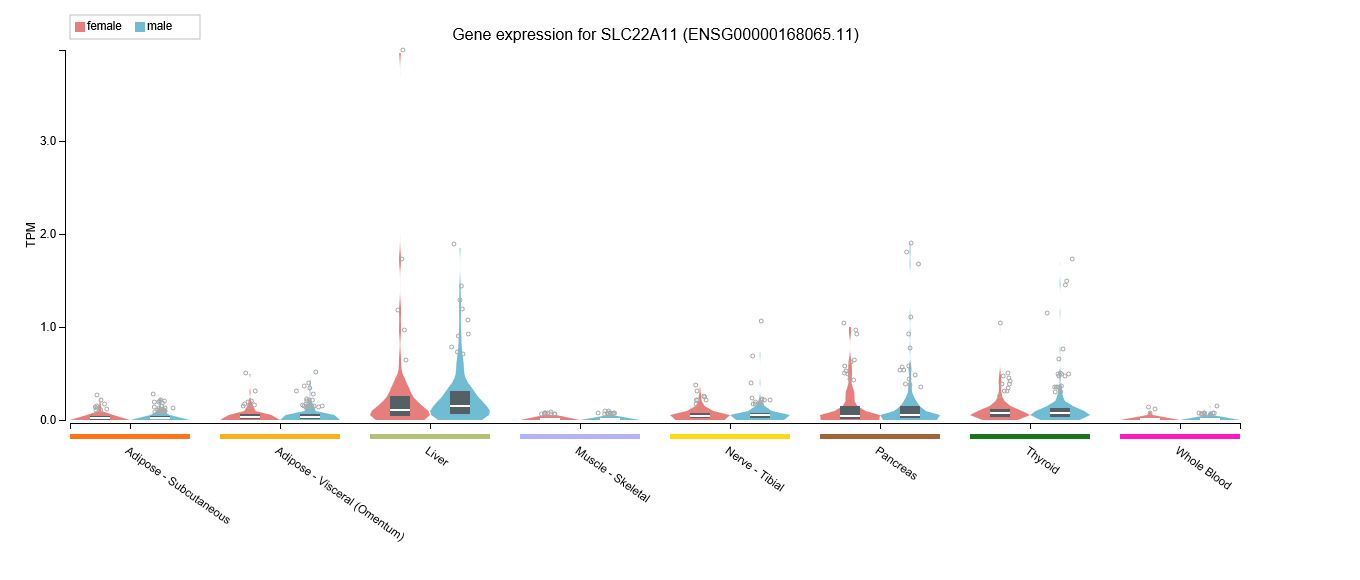

c) ***ZNF45***

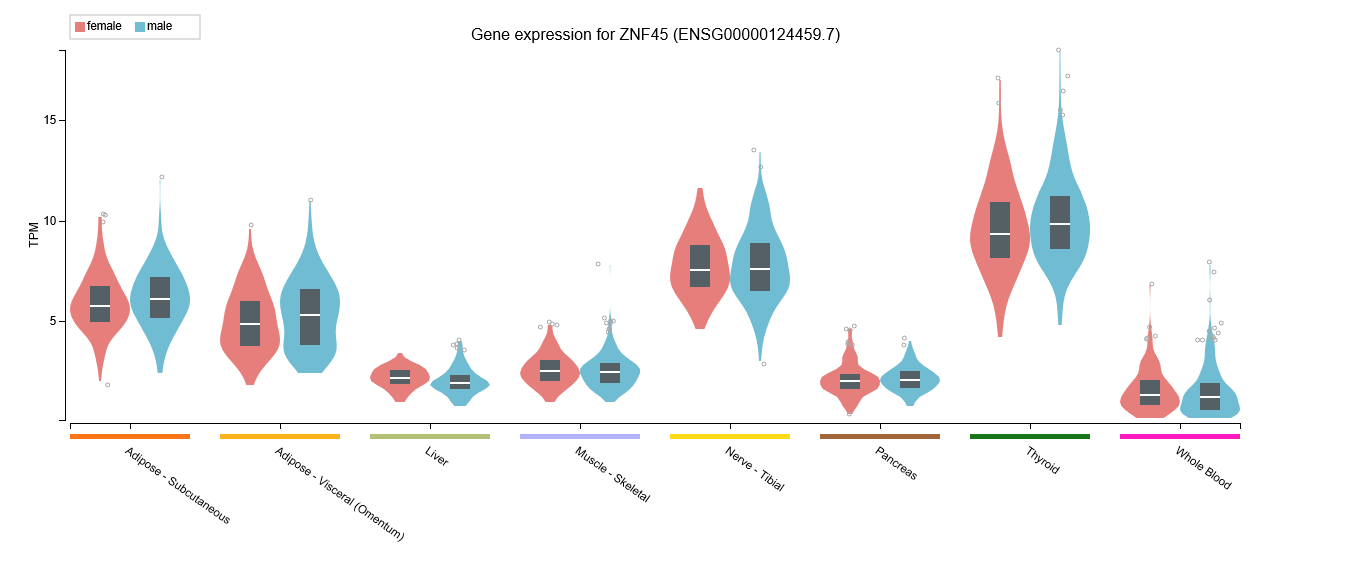

**Fig.S10** Gene expression of *BAI3, SLC22A11* and *ZNF45* (mRNA levels) [GTEx portal]

**Fig.S11 Gene regulatory features of *BAI3* locus**

a) **ATAC-Seq, DNase I peaks and regulatory histone marks aroud rs6913677 in human subcutaneous abdominal adipose tissue**

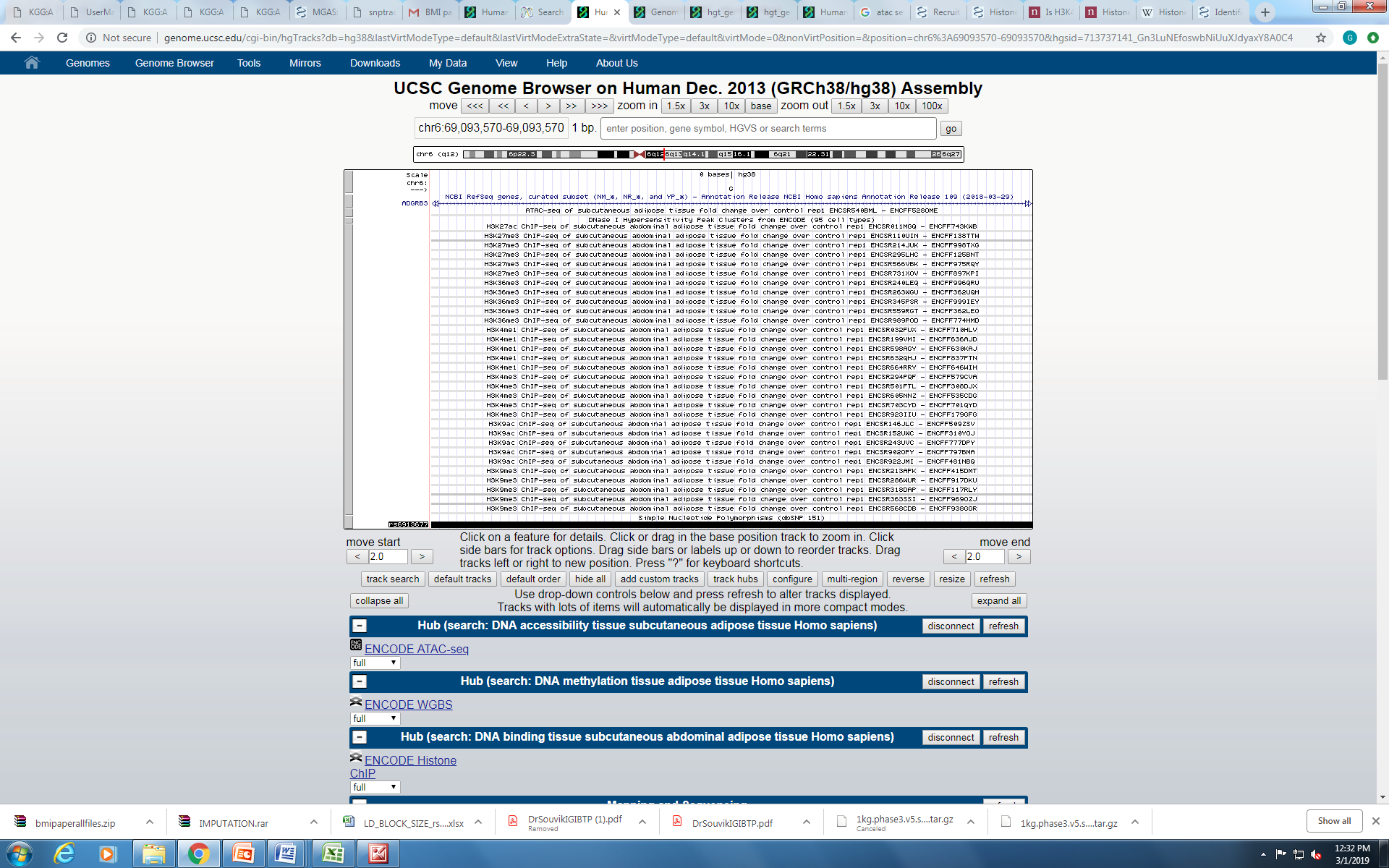

**Fig.S11 (a)** ATAC-seq and DNase I peaks: open chromatin region. Active histone marks for gene transcription: H2K27ac, H3K4me1, H3K9ac, H3K4me3, and H3K36me3. Repressive histone mark: H3K27me3 and H3K9me3 [ENCODE]

b) **Chromatin interaction potential, transcription factor binding and DNA methylation status around rs6913677 variant region in human adipose tissue**

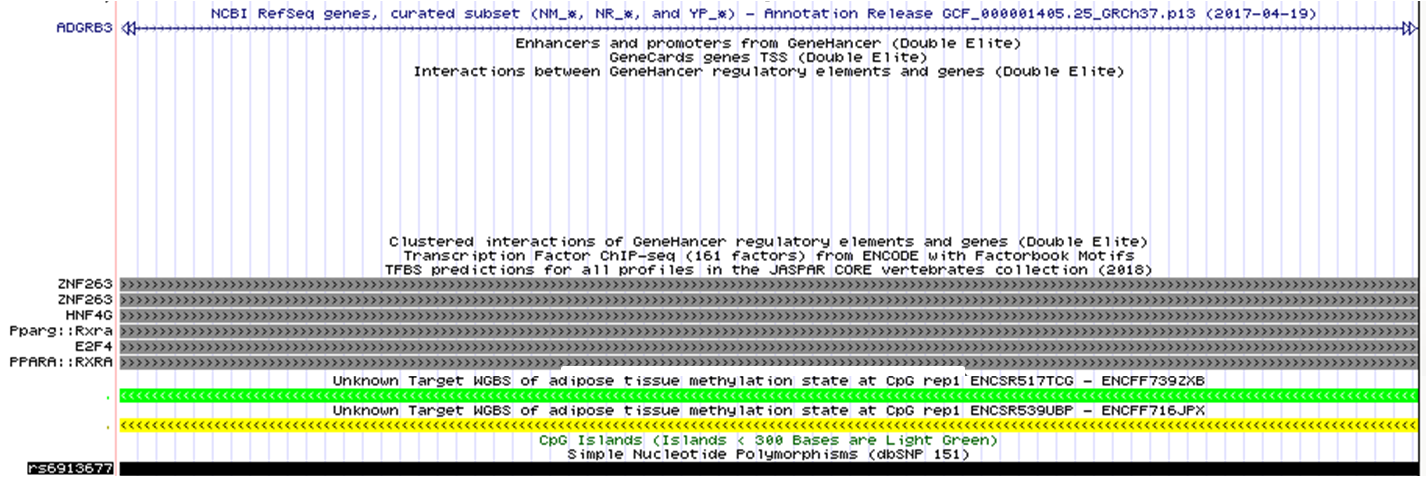

**Fig.S11 (b)** Chromatin interaction potential data was derived from GeneHancer database. Predicted and experimentally recognized transcription factors binding to variant were retrieved from JASPAR and ENCODE database where black color signify strongest binding and light gray as fragile binding. Whole Genome Bisulphite Sequencing data (WGBS) of adipose tissue was acquired from ENCODE browser. Color representation: red (100% of sequenced reads are methylated), yellow (50% of sequenced reads are methylated), green (0% of sequenced reads are methylated)

**c)** **Conservation of predicted TF motifs at rs6913677 variant region**

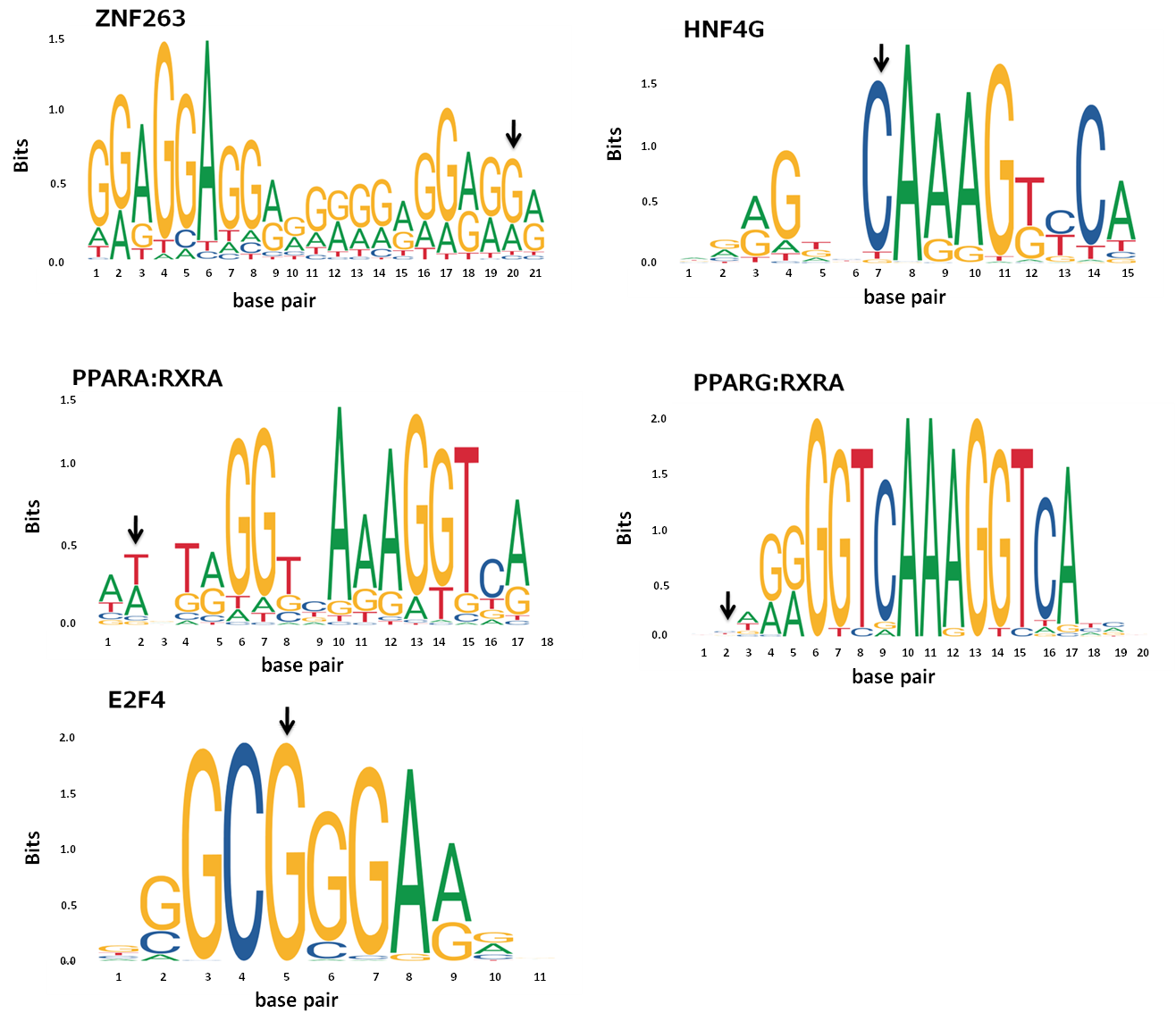

**Fig.S11(c)** DNA binding motifs at variant region for predicted transcription factors were explored from JASPAR database. Arrow indicate position of variant

**Fig.S12 Gene regulatory features of *SLC22A11* locus**

a) **ATAC-Seq, DNase I peaks and regulatory histone marks aroud rs2078267 in human subcutaneous abdominal adipose tissue**

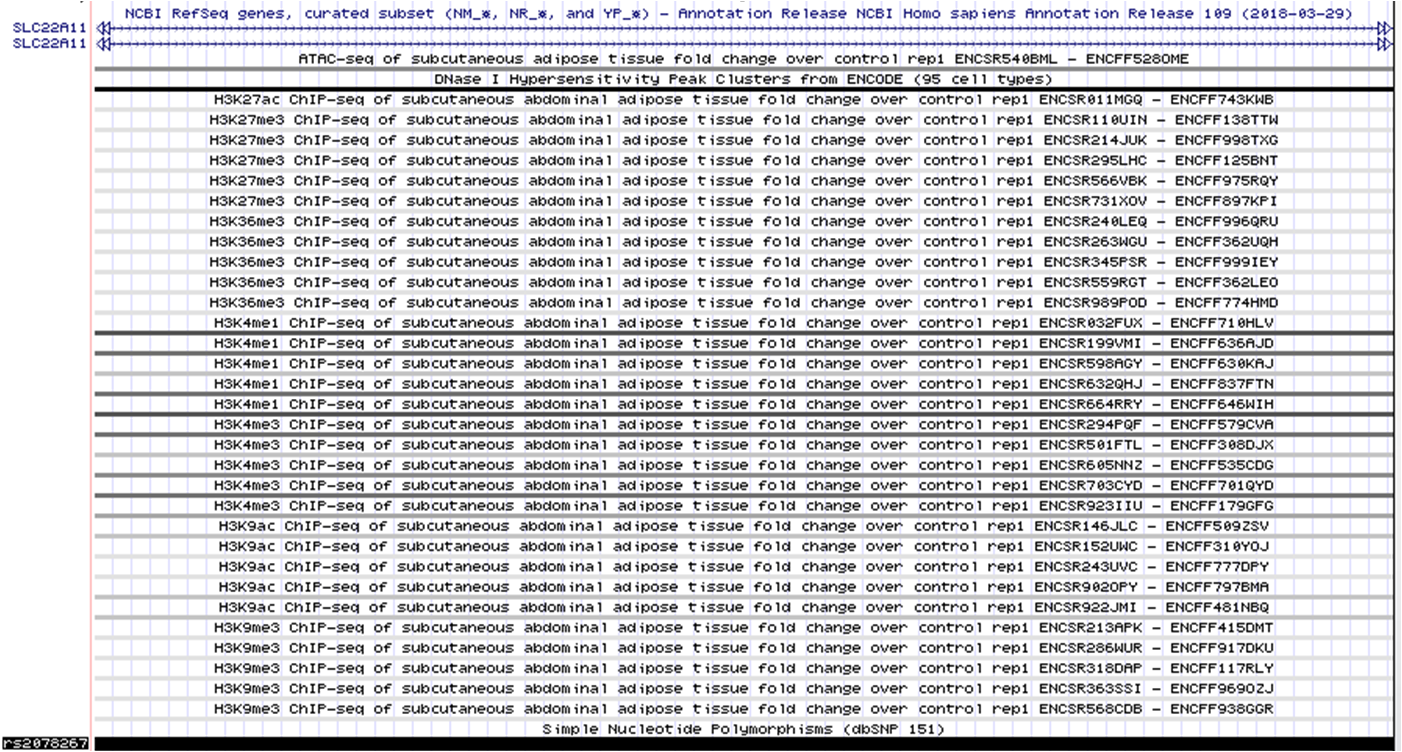

**Fig.S12 (a)** ATAC-seq and DNase I peaks: open chromatin region. Active histone marks for gene transcription: H2K27ac, H3K4me1, H3K9ac, H3K4me3, and H3K36me3. Repressive histone mark: H3K27me3 and H3K9me3 [ENCODE]

**b) Chromatin interaction potential, transcription factor binding and DNA methylation status around rs2078267 variant region in human adipose tissue**

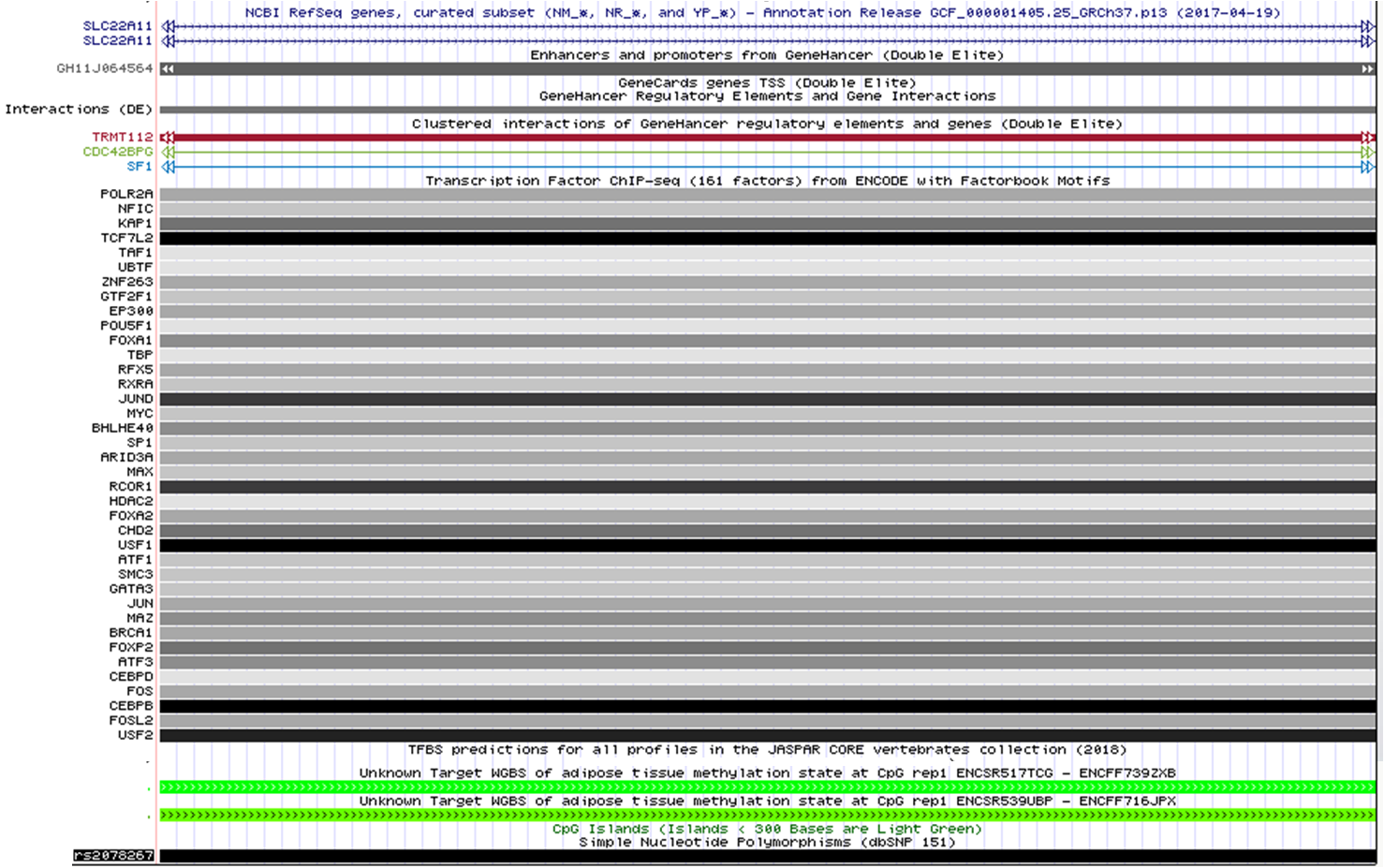

**Fig.S12 (b)** Chromatin interaction potential data was derived from GeneHancer database. Predicted and experimentally recognized transcription factors binding to variant were retrieved from JASPAR and ENCODE database where black color signify strongest binding and light gray as fragile binding. Whole Genome Bisulphite Sequencing data (WGBS) of adipose tissue was acquired from ENCODE browser. Color representation: red (100% of sequenced reads are methylated), yellow (50% of sequenced reads are methylated), green (0% of sequenced reads are methylated)

**Fig.S13 Gene regulatory features of *ZNF45* locus**

**a) ATAC-Seq, DNase I peaks and regulatory histone marks aroud rs8100011 in human subcutaneous abdominal adipose tissue**

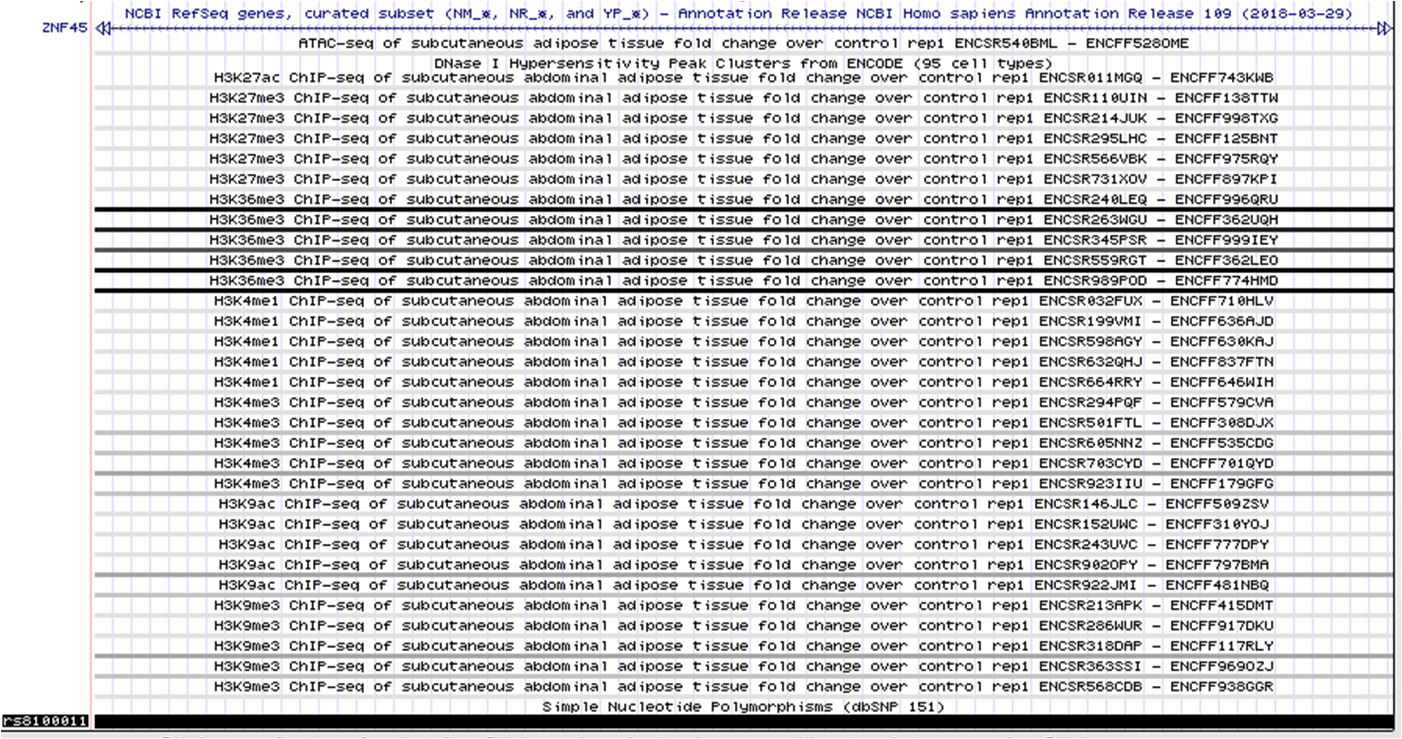

**Fig.S13 (a)** ATAC-seq and DNase I peaks: open chromatin region. Active histone marks for gene transcription: H2K27ac, H3K4me1, H3K9ac, H3K4me3, and H3K36me3. Repressive histone mark: H3K27me3 and H3K9me3 [ENCODE]

***b)* Chromatin interaction potential, transcription factor binding and DNA methylation status around rs8100011 variant region in human adipose tissue**

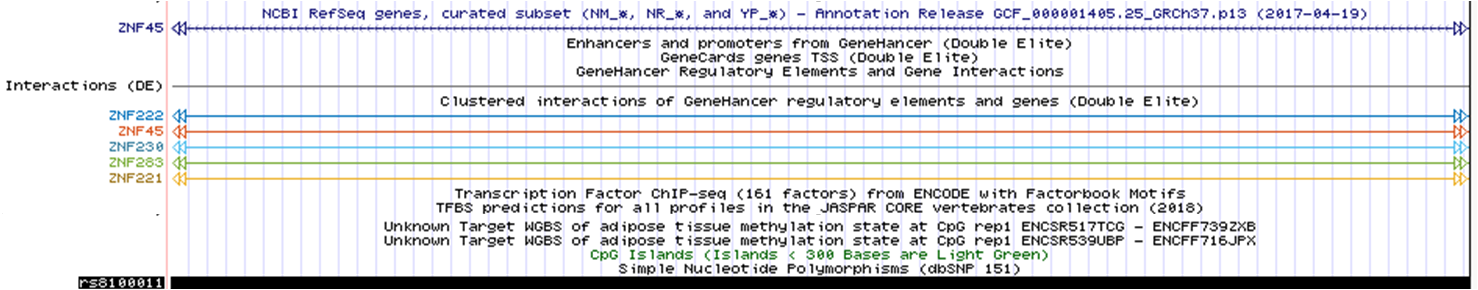

**Fig.S13 (b)** Chromatin interaction potential data was derived from GeneHancer database. Experimentally recognized transcription factors binding to variant were retrieved from ENCODE and JASPAR database where black color signify strongest binding and light gray as fragile binding. Whole Genome Bisulphite Sequencing data (WGBS) of adipose tissue was acquired from ENCODE browser. Color representation: red (100% of sequenced reads are methylated), yellow (50% of sequenced reads are methylated), green (0% of sequenced reads are methylated)

**Fig.S14 Gene expression profiles of *BAI3, SLC22A11* and *ZNF45* in human blood**

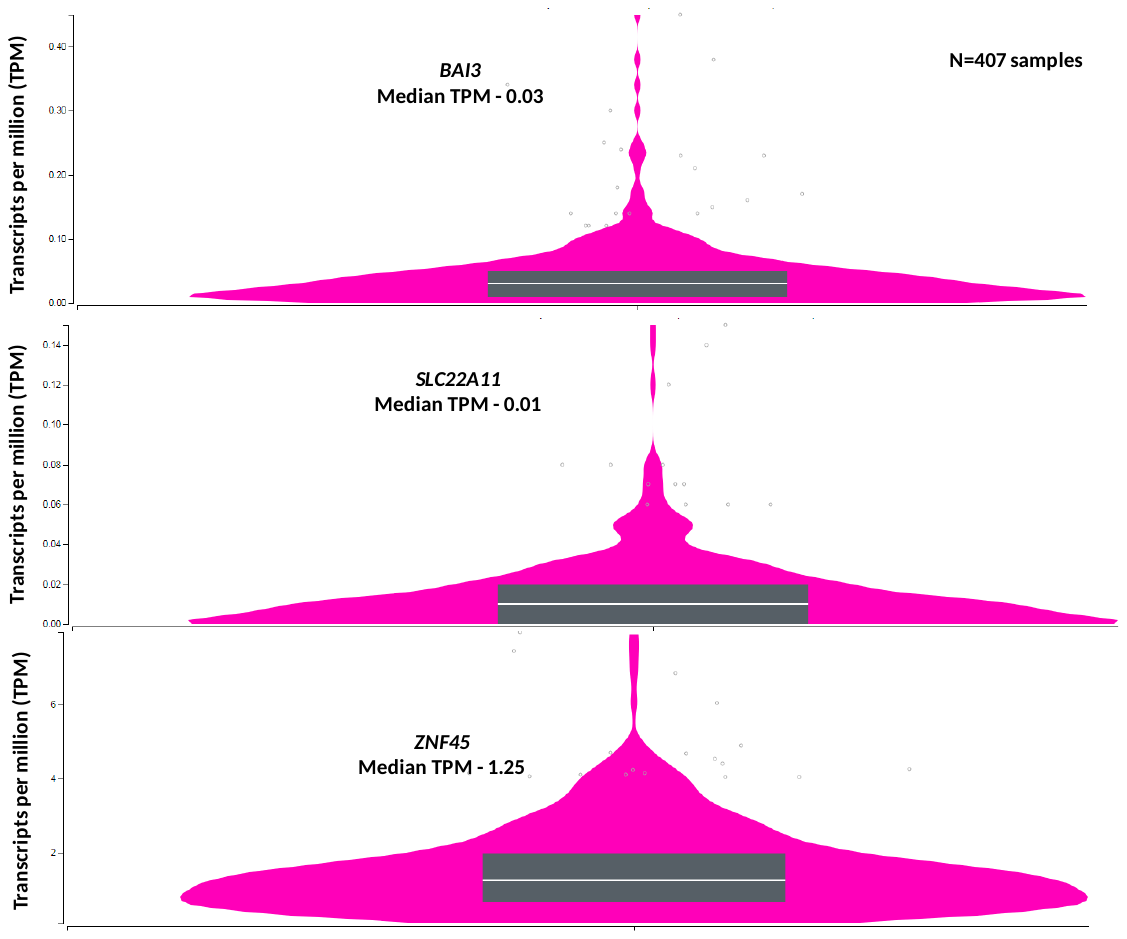

**Fig.S14**

Data was derived from GTEx portal. Number of tissues samples for whole blood - 407

**Fig.S15 DNA methylation marks around *BAI3, SLC22A11* and *ZNF45* loci in CD14 (positive) monocytes, B-cells and T-cells**

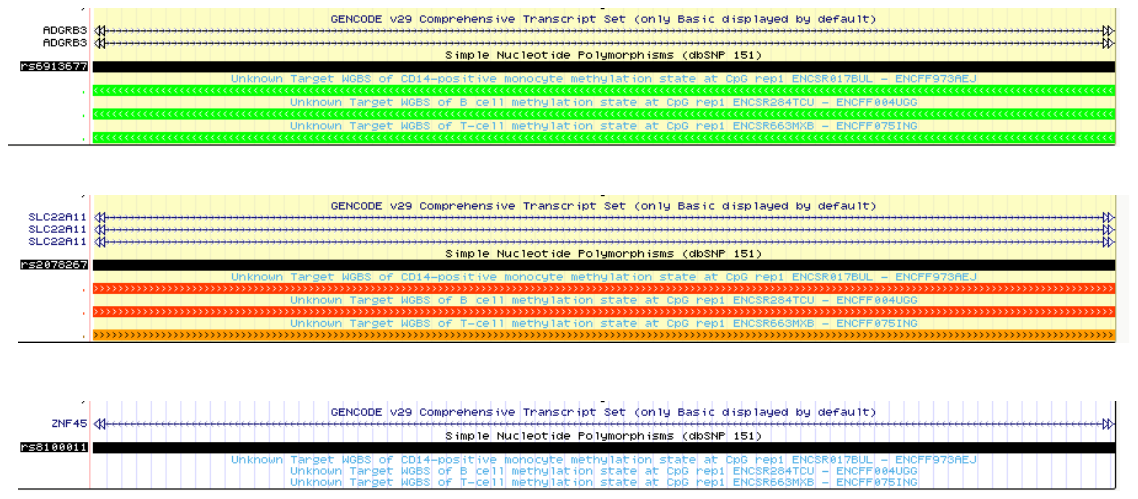

**Fig.S15** Whole Genome Bisulphite Sequencing data (WGBS) was acquired for CD14 (positive) monocytes, B-cells and T-cells for *BAI3, SLC22A11* and *ZNF45* variants from ENCODE browser. Color representation: red (100% of sequenced reads are methylated), yellow (50% of sequenced reads are methylated), green (0% of sequenced reads are methylated)
